# Reference-free multiplexed single-cell sequencing identifies genetic modifiers of the human immune response

**DOI:** 10.1101/2023.05.29.542756

**Authors:** George C. Hartoularos, Yichen Si, Fan Zhang, Pooja Kathail, David S. Lee, Anton Ogorodnikov, Yang Sun, Yun S. Song, Hyun Min Kang, Chun Jimmie Ye

**Author notes:** These authors contributed equally (Yichen Si, Fan Zhang).

## Abstract

Multiplexed single-cell sequencing (mux-seq) using single-nucleotide polymorphisms (SNPs) has emerged as an efficient approach to perform expression quantitative trait loci (eQTL) studies that map interactions between genetic variants and cell types, cell states, or experimental perturbations. Here we introduce the *clue* framework, a novel approach to encode mux-seq experiments that eliminates the need for reference genotypes and experimental barcoding. The *clue* framework is made possible by the development of *freemuxlet*, an algorithm that clusters cells based on SNPs called from single-cell RNA-seq or ATAC-seq data. To demonstrate the feasibility of *clue*, we profiled the surface protein and RNA abundances of peripheral blood mononuclear cells from 64 individuals, stimulated with 5 distinct extracellular stimuli — all within a single day. Our analysis of the demultiplexed data identified rare immune cell types and cell type-specific responses to interferon and toll-like receptor stimulation. Furthermore, by integrating genotyping data, we mapped response eQTLs specific to certain cell types. These findings showcase the potential and scalability of the *clue* framework for reference-free multiplexed single-cell sequencing studies.

## Introduction

Understanding the genetic architecture of gene expression remains a critical challenge in human genetics. The overwhelming enrichment of disease-associated variants in the cis-regulatory regions of the genome points to the crucial role of transcription regulation in conferring disease risk^1, 2^. Although expression quantitative trait loci (eQTL) studies in bulk tissues have identified numerous genetic variants associated with proximal gene expression, their enrichment for disease-associated variants remains modest^3, 4^. This might be because disease-causing variants affect enhancer rather than promoter activities, modifying gene expression in particular cell types, cell states, or in response to specific environmental factors. In such situations, it can be challenging to identify eQTLs that interact with cellular states using bulk gene expression analysis, as the composition of cell types and the molecular states of cells within the same type may vary between individuals, and functionally important cell populations could be rare^5^. One method for mapping eQTL interactions is to sort and perturb specific cell types and then profile their gene expression. However, this approach is cost prohibitive for large population cohorts, can be susceptible to experimental confounding, and fails to capture heterogeneity within sorted populations. Consequently, there is a need for more efficient and unbiased methods for mapping eQTL interactions in the human genome.

Multiplexed single-cell sequencing (mux-seq) using single-nucleotide polymorphisms (SNPs) as sample barcodes has enabled population-scaled studies for assessing the impact of case-control status^6^, experimental perturbations^7^, and genetic variants on gene expression across single cells^8^. Recently, our analyses of mux-seq data revealed that cell type-specific *cis*-eQTLs are more enriched for disease associations than those shared across circulating immune cell types^6^. Mux-seq is highly adaptable, requires minimal experimental modification over standard single-cell sequencing workflows, and has been shown to be compatible with single-cell RNA-seq, single-nuclei RNA-seq^9^, and CITE-seq^10^. However, current mux-seq implementations require either reference genotypes or experimental barcoding to unambiguously assign cells to each sample. This limitation precludes the application of mux-seq for studies involving cells that are sensitive to manipulation or for samples where genotyping may not be feasible due to privacy or availability concerns.

Here, we introduce *clue*, a framework for mux-seq experiments that eliminates the need for reference genotypes or experimental barcoding. *Clue* incorporates a series of pooling schemes for efficient experiment encoding and a demultiplexing algorithm to determine the unique sample identity of each cell. This is made possible by the development of *freemuxlet*, an extension of demuxlet^11^ that allows clustering of genetically-identical cells from pooled scRNA- and scATAC-seq experiments without reference genotypes. We applied *clue* to investigate the response of peripheral blood mononuclear cells (PBMCs) to five different agonists targeting the type I and type II interferon responses (recombinant IFNβ and IFNγ), viral sensing (R848), inflammatory response (TNFɑ), and broad immune cell activation (PMA/I). The *clue* framework allowed us to perform multiplexed CITE-seq across 384 samples from 64 individuals across 12 pools in just one day. Analyzing 134,831 cells, we discovered rare cell types and identified cell type-specific transcriptional responses that were validated by bulk RNA-sequencing. We identified shared and specific transcriptional responses to interferons in monocytes, highlighted by the discovery of specific effects in non-classical monocytes related to a migratory phenotype induced by type I interferon and complement activation induced by type II interferon. Lastly, by integrating imputed genotyping data, we mapped cell type-specific cis response eQTLs (cis-reQTLs) to each stimulation, identifying specific associations in R848-stimulated naive B cells (*IFITM2*) and IFNβ-stimulated classical monocytes (*UBE2F*). These findings showcase the efficiency and robustness of *clue* as a framework for reference-free multiplexed single-cell sequencing.

## Results

### *clue*: genetic multiplexing without reference genotypes

Here, we introduce *clue* (<u>c</u>ompressed, lossless, <u>u</u>nambiguous multipl<u>e</u>xing), a workflow for multiplexed single-cell sequencing (mux-seq) that enables population-scale single-cell studies without reference genotypes or experimental barcoding (**Fig. 1A**). We illustrate the key features of *clue* utilizing a toy study that profiles *n* individuals over *r* conditions, where *r* < *n*. The conditions could be different perturbations (as illustrated), time points, or aliquots of the same cells. The core of *clue* is a *p* × *n* pooling matrix that assigns each of *n* samples to one of *p* pools. After single-cell profiling of the pools, the resulting data is first analyzed through *freemuxlet*, a novel algorithm that clusters cells based on genetic variants identified directly from the single-cell sequencing data. Genetic clusters of cells from different pools are then demultiplexed, where each cell is correctly assigned to an individual and condition.

**Figure 1.**
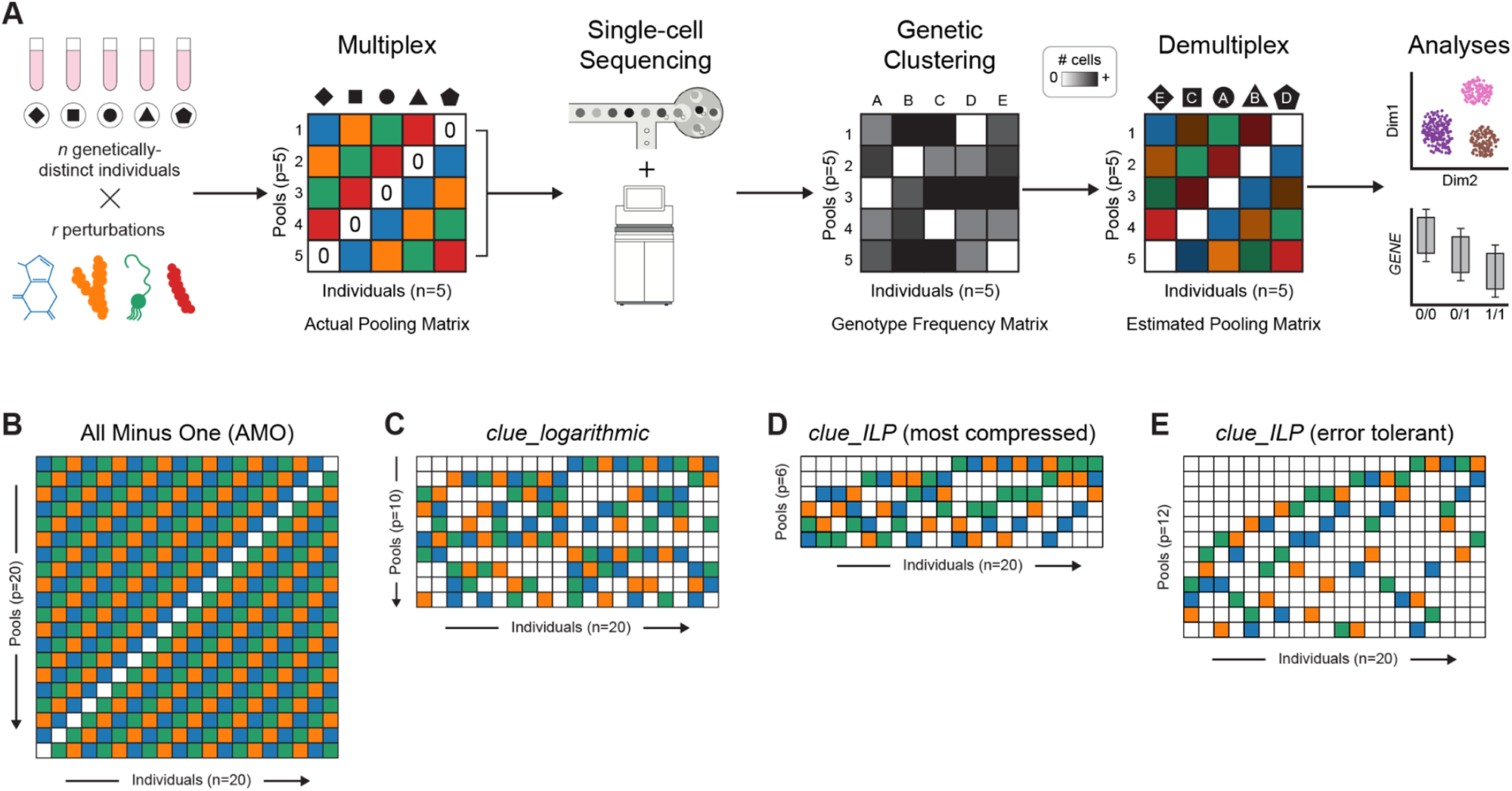
Overview of the *clue* framework. **A**, Illustrative schematic of the *clue* framework using the all-minus-one (AMO) pooling matrix, in which cells from one individual are omitted per pool. After single-cell sequencing, cells are genetically clustered and can be demultiplexed by identifying which samples are absent in each pool. Off-diagonal variance in cell numbers in the genotype frequency matrix is due to technical variability (e.g. unequal mixing of cells). The estimated pooling matrix is overlaid with the shading from the genotype frequency matrix to indicate the number of cells observed per individual-pool. **B**, For a toy example of 20 individuals and 3 perturbations, an AMO pooling matrix is identifiable but not most compact. **C**, *clue_logarthmic* is a more compressed pooling matrix with fewer pools. *clue_ILP* enables discovery of **D**, optimal (i.e. most compressed) pooling schemes and **E**, those that are error tolerant and batch effects minimized.

In order to ensure successful demultiplexing, *clue* aims to produce a pooling matrix that assigns the *n* × *r* samples to a minimum number of pools while meeting three key objectives:

- Identifiability: each cell can be uniquely assigned to a sample (e.g., individual and condition);
- Robustness: samples are distinguishable while tolerating errors in the pooling or genetic clustering;
- Balance: cells from each individual and each condition are uniformly distributed across pools.

There are several different multiplexing schemes that can achieve these objectives. The naive all-minus-one (AMO) scheme which omits each individual’s cells from exactly one pool meets the identifiability objective but requires *n* pools, which limits the experimental efficiency of sample multiplexing (**Fig. 1B**). The *clue_logarithmic* scheme assigns samples using at least *p* = 2 × *log*_2_(*n*) pools motivated by previous work describing logarithmic encoding^12^, which achieves significant compression compared to AMO and is experimentally easy to perform (**Fig. 1C**). In a toy example, multiplexing *n* = 20 individuals over *r* = 3 conditions can be encoded using *p* = 10 pools. However, it may not be the most compressed or error-tolerant scheme.

The *clue_ILP* scheme uses integer linear programming (ILP) to identify the optimal multiplexing scheme (Methods). This scheme can further be optimized for condition randomization and error tolerance, by distributing the samples and maximizing the differences in the multiplexing matrix profiles, respectively (**Fig. 1E**, **Fig. S1**). In our toy example, the most compressed scheme only needed *p* = 6 pools to ensure demultiplexing (**Fig. 1D**), and an error-tolerant multiplexing scheme required *p* = 12 (**Fig. 1E**).

### freemuxlet: genetic clustering of single cells without reference genotypes

The *clue* framework requires the ability to group genetically identical cells without relying on reference genotypes obtained from a genotyping array or sequencing. To meet this need, we developed *freemuxlet*, an approach based on demuxlet^11^ that genetically clusters cells using only SNPs captured from multiplexed single-cell sequencing data (**Fig. 2A**). Instead of relying on reference genotypes, freemuxlet uses unsupervised learning to efficiently cluster genetically-identical cells and identify heterotypic multiplets — droplets containing two or more cells from different individuals.

**Figure 2.**
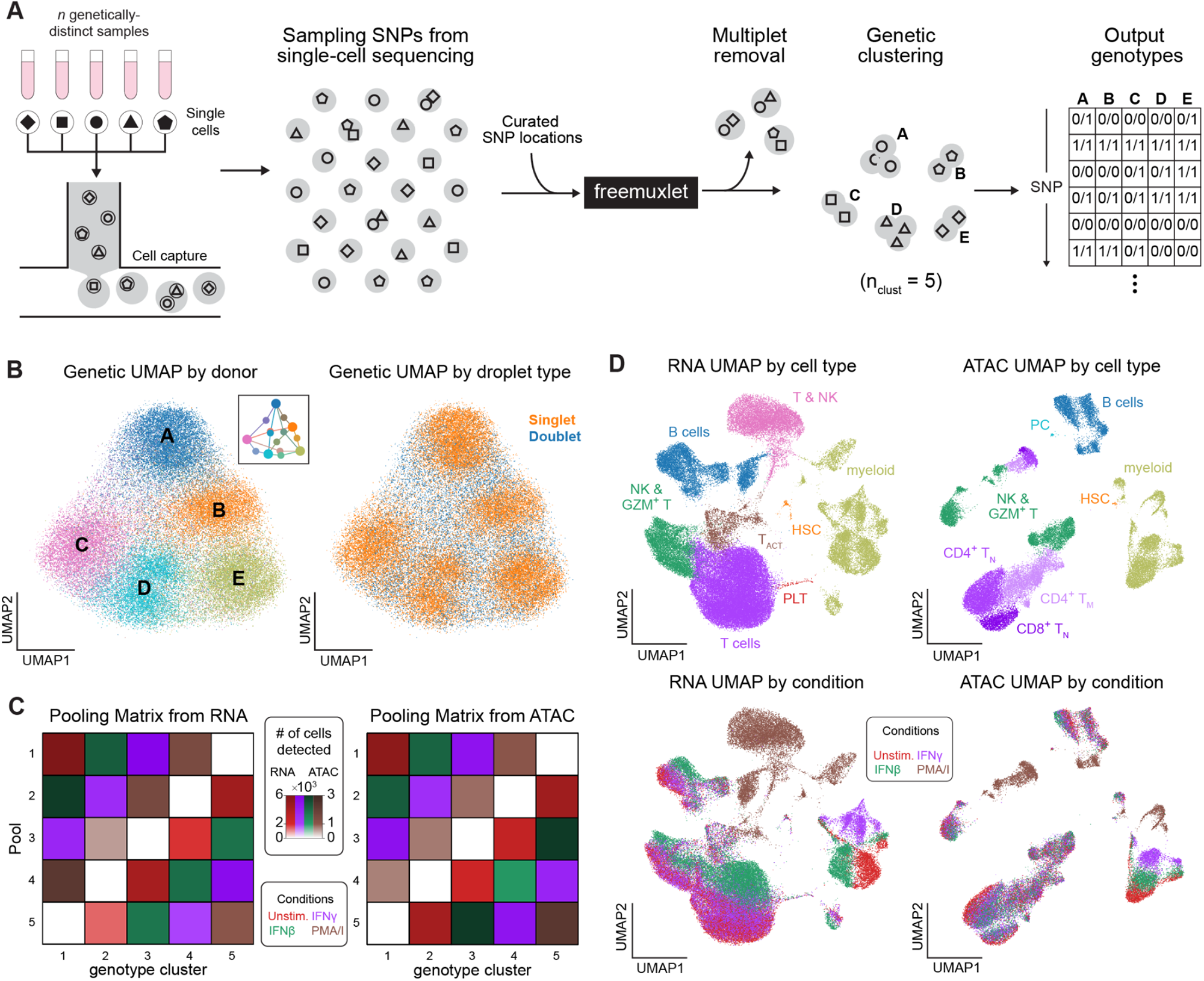
Overview of freemuxlet as applied to *clue* data. **A**, Schematic of the freemuxlet algorithm, in which single-cell sequencing data and a curated set of loci are input, and genetically-distinct clusters of singlets and a variant calling format (VCF) genotype file are output. **B**, Visualizing the pairwise genetic distance between droplets in UMAP space shows 5 distinct clusters corresponding to the 5 input individuals, as well as putative doublets that embed between constituent donor clusters. **C**, The estimated pooling matrix of singlets from the AMO experiment recapitulates the actual pooling matrix for both RNA and ATAC assays. Stimulation conditions are introduced to take advantage of redundancy. **D**, The resulting single-cell transcriptome and chromatin accessibility profiles visualized in UMAP space show heterogeneity due to both cell type and stimulation condition.

At its core, *freemuxlet* uses a modified Expectation-Maximization (E-M) algorithm to assign barcoded droplets containing cells to clusters, updating the cluster assignments iteratively. A droplet is labeled as a singlet if it has been successfully assigned to a single cluster, or a multiplet if it cannot be unequivocally assigned to any given cluster. Compared to existing genetic clustering algorithms like scSplit^13^, vireo^14^, and souporcell^15^, freemuxlet stands out with two key features. Firstly, freemuxlet incorporates a singlet score based solely on allele frequencies, significantly improving the quality of initial clustering and the speed and accuracy of convergence. This becomes especially crucial when dealing with a large number of multiplexed individuals or high multiplet rates. Secondly, freemuxlet refines cluster assignments using an identity-aware Bayes factor that leverages both base and read quality to extract the maximum information from the sequence data. Indeed, these two aspects may explain the superior performance of *freemuxlet* compared to existing methods^16^.

To showcase the performance of *freemuxlet* and its suitability for the *clue* framework, we conducted multiplexed single-cell RNA- and ATAC-seq experiments assaying PBMCs from 5 individuals across 4 conditions using the AMO multiplexing scheme. By using a set of curated SNP locations (Methods), *freemuxlet* was able to group cells based on their genotypes estimated from either the single-cell RNA-seq or the ATAC-seq data. The results from the ATAC-seq data, visualized using Uniformed Manifold Approximation and Projection (UMAP) of the pairwise genetic distances, showed 5 distinct clusters of singlets and putative doublets occupying regions of the UMAP between clusters (**Fig. 2B, Fig. S2A**). Analysis of the RNA-seq data revealed allele-specific expression only in certain cell types or in response to certain perturbations, which highlights the importance of incorporating allele frequency in the clustering algorithm (**Fig. S2B**). The demultiplexing results from both the RNA-seq and ATAC-seq data matched the pooling matrix (**Fig. 2C**) and were consistent with demultiplexing using demuxlet with reference genotypes (**Fig. S2C**). Furthermore, the genotypes detected from both RNA-seq and ATAC-seq were in agreement with those obtained from a SNP genotyping array (**Fig. S2D**). By visualizing the resulting demultiplexed single-cell RNA- and ATAC-seq profiles using UMAP, we observed cells clustered primarily by type, and to a lesser extent by stimulation. Differential expression analysis of the same cell type between different conditions provides further evidence of correct demultiplexing (**Fig. 2D**, **Fig. S3**). For example, PMA/I stimulation induced the strongest effects, with stimulated cells of each major cell type forming distinct clusters from unstimulated cells of the same type. On the other hand, IFNγ stimulation had the weakest effects, with stimulated cells mostly clustering with unstimulated cells. These results show that *freemuxlet* is a reference-free method for clustering cells based on genetic variation, suitable for both single-cell RNA-seq and ATAC-seq data and can be deployed in the *clue* framework to enable population-scale single-cell sequencing studies.

### Application of *clue* to parse cell type-specific immune responses

To demonstrate the suitability and scalability of the *clue* framework for population-scale single-cell sequencing studies, we performed a multiplexed single-cell CITE-seq experiment to study the genetic modulation of immune response in PBMCs. We assayed PBMCs from 64 female, non-hispanic white healthy individuals either at rest (unstimulated control) or stimulated with one of five immunomodulatory molecules: tumor necrosis factor alpha (TNFα), interferons gamma (IFNγ) and beta (IFNβ), TLR7/8 agonist resiquimod-848 (R848), and phorbol-myristate-acetate with ionomycin (PMA/I) (**Fig. 3A**). The cells were profiled at 9 hours post-stimulation, a time point that was found to induce potent transcriptional effects in response to most stimuli from bulk RNA-sequencing of PBMCs (**Fig. S4A–B**). The full experiment of 384 samples (64 individuals by 6 conditions) was profiled in 12 pools according to a pooling matrix produced by *clue_logarithmic*. The matrix assigned 32 genetically-distinct samples per pool, utilizing an internally-symmetric tree structure that is experimentally simple to execute (**Fig. 3B**). Upon sequencing, alignment, genetic clustering of cells using *freemuxlet*, and demultiplexing, we correctly reconstructed 98.9% elements of the pooling matrix (760/768 matrix elements; **Fig. 3C**, **Fig. S5A–B**). The errors were due to a mis-pooling event (genotype cluster 11) and the loss of one individual’s cells during culture due to low viability (genotype cluster 59; **Fig. S5C**). Although not explicitly optimized to be error-tolerant, the multiplexing scheme was robust to these errors and cells were assigned to 64 individuals across 6 conditions.

**Figure 3.**
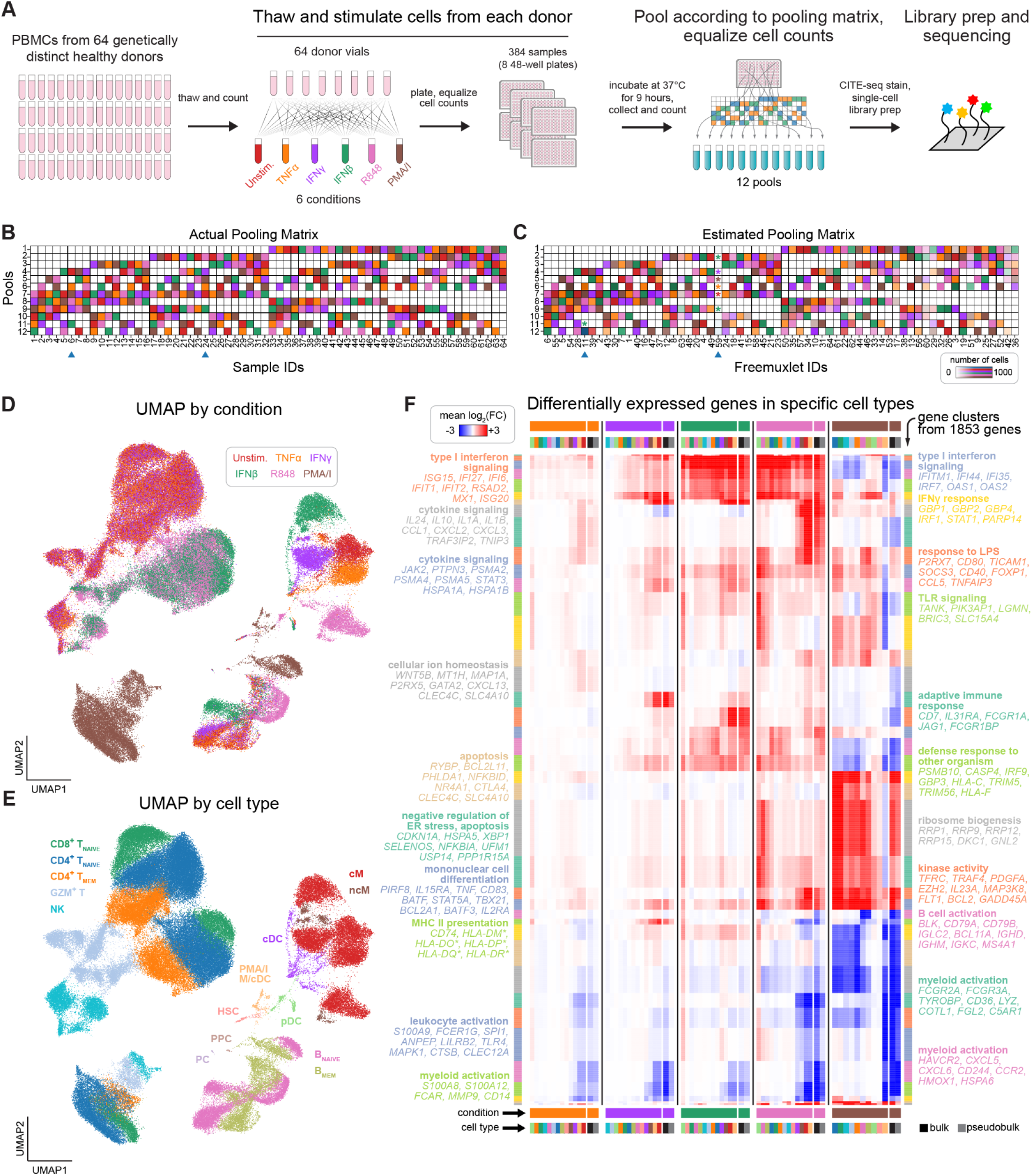
The *clue* framework enables single-cell profiling of 384 samples in 12 reactions. **A**, Experimental overview. PBMCs from 64 donors were incubated with 5 immunomodulatory stimulants for 9 hours, then pooled and sequenced. **B–C**, The actual pooling matrix and estimated pooling matrix from *freemuxlet* show near-perfect concordance. Two deviations (blue arrows), one mis-pooling event (genotype cluster 11) and one instance of cell loss (low recovery of a low viability sample, genotype cluster 59), are highlighted with asterisks. Demultiplexing was robust to these errors. **D–E**, Dimensionality reduction with UMAP and clustering with Leiden shows heterogeneity in gene expression from both stimulation condition (**D**) and cell type (**E**). **F**, Heatmap of differentially expressed genes comparing stimulation conditions to controls in each cell type. Genes are *k*-means clustered to yield gene modules with significant functional enrichment in immune-relevant biological pathways. Pseudobulks across all cell types per condition are concordant with bulk RNA data.

The demultiplexed CITE-seq data was visualized with UMAP, and the cell clusters determined by Leiden clustering generally tracked with cell type and stimulation and not with batch or other technical parameters (**Fig. 3D–E**; **Fig. S6A–C**). T and NK cells stimulated by IFNγ and TNFα clustered together with control cells and separately from those stimulated by IFNβ and R848. For B cells, R848- and IFNβ-stimulated cells clustered together, whereas IFNγ-stimulated and control cells clustered together. In monocytes, cells stimulated by each stimulus formed their own distinct cluster. PMA/I-stimulated lymphoid cells clustered out separately from other stimuli, replicating the strong effects observed in the AMO and bulk experiments, while PMA/I-stimulated myeloid cells were significantly depleted, likely due to differentiation and adhesion to the tissue culture plate after stimulation (Methods).

After performing differential expression (DE) analysis between stimulated and unstimulated cells, we identified 1853 DE genes in at least one cell type and one perturbation (*log_2_(Fold Change)* > 1, *p*_adj_ < 0.05). We then used K-means clustering to group these genes into functional modules that were enriched for immune-related pathways such as cytokine signaling, activation, response to exogenous stimulation (e.g. LPS, virus, other organism), type I IFN signaling, adaptive immune response, and apoptosis (**Fig. 3F**, **Fig. S6D–G**, **Table S1**). TNFα induced the lowest fold change, except for genes related to cellular ion homeostasis (e.g., *MT1*), while PMA/I induced the highest fold change, especially for genes related to ribosome biogenesis, RNA processing, and proliferation. IFNγ, IFNβ, and R848 induced intermediate fold changes for genes implicated in TLR signaling, defense response, and antigen processing/presentation. Importantly, the log fold change estimates from the pseudobulk analysis of the scRNA-seq data were highly consistent with those estimated from the bulk PBMC RNA-sequencing data after 9 hours of stimulation (**Fig. 3F**, **Fig. S4C**). These findings demonstrate the *clue* framework can be deployed at scale to map cell type-specific responses to immune modulation in circulating immune cells.

### Identification of rare lymphoid cell types and stimulation-specific transcriptional responses

To assess the impact of stimulation on PBMC subsets, we next analyzed the data after subclustering cells based on their lineage (Methods). We first jointly analyzed T and NK cells, identifying 22 distinct cell clusters consisting of naive and memory T cell subsets, gamma delta T cells (T_ɣδ_), mucosal associated invariant T (MAIT) cells, and NK cells (**Fig. 4A–B**). Within naive CD4^+^ and CD8^+^ T cells (confirmed by CD45RA^+^ surface expression), we identified 4 subclusters that were differentiated by the expression of *SELL* (CD62L) and *CD69* (CD69) transcript and protein, indicating a spectrum of stimulation-specific phenotypes. Cluster 7 consisted of R848-stimulated CD4^+^ and CD8^+^ cells, which suggested condition-specific effects shared between the T cell subsets. Activated (CD45RO^+^, cluster 5) and resting (CD45RA^+^, cluster 6) Tregs were marked by their specific expression of *FOXP3*. Among other CD45RO^+^ CD4^+^ T cells, we identified T_h_2 cells (*CDO1*, *PTGDR2*; cluster 10) and a cluster of cells that did not polarize to any particular T helper cell state (*CXCR3*, *CXCR5*, *RORC*, *CCR4*, *CCR5*, *CCR6*; cluster 9; **Fig. S7A**). Notably, we found a subset of CD8^+^ T cells with high transcript and protein expression of *ITGAE* (CD103) (cluster 11), which is a marker for tissue resident memory cells (T_RM_). Among the cytotoxic cells marked by the expression of granzyme family members (GZM*^+^*), we identified expected subsets of memory CD8^+^ T cells, T_ɣδ_ cells, MAIT cells, and NK cells. We also found a cluster of CD56-expressing cells with high expression of *IL2RA* (CD25) and c-kit (CD117), and lower expression of granzymes and transcription factors (TFs) *EOMES* and *TBX21* (Tbet), supporting their annotation as circulating innate lymphoid cells (ILCs)^17, 18^ (**Fig. S7B**). Lastly, we identified two small populations (clusters 13 and 14) marked by the expression of TFs *ZNF683* (HOBIT) and *IKZF2* (HELIOS) and differentiated by the expression of *MME* (CD10) (**Fig. S7C–D**). Cluster 13 is labeled as immature T cells or common lymphoid progenitors (CLPs)^19, 20^, an annotation further supported by their expression of other genes shown to be involved in T cell development (e.g. *SOX4*^21^*, FXDY2*^22^; shared with *SELL*^+^ and *SELL*^int^ naive subsets, respectively; **Fig. S7E**). Cluster 14 resembles the recently-described HOBIT^+^/HELIOS^+^ T cells^23^, an unexpected finding in circulation since HOBIT has been shown to identify non-circulating resident memory T cell precursors^24^.

**Figure 4.**
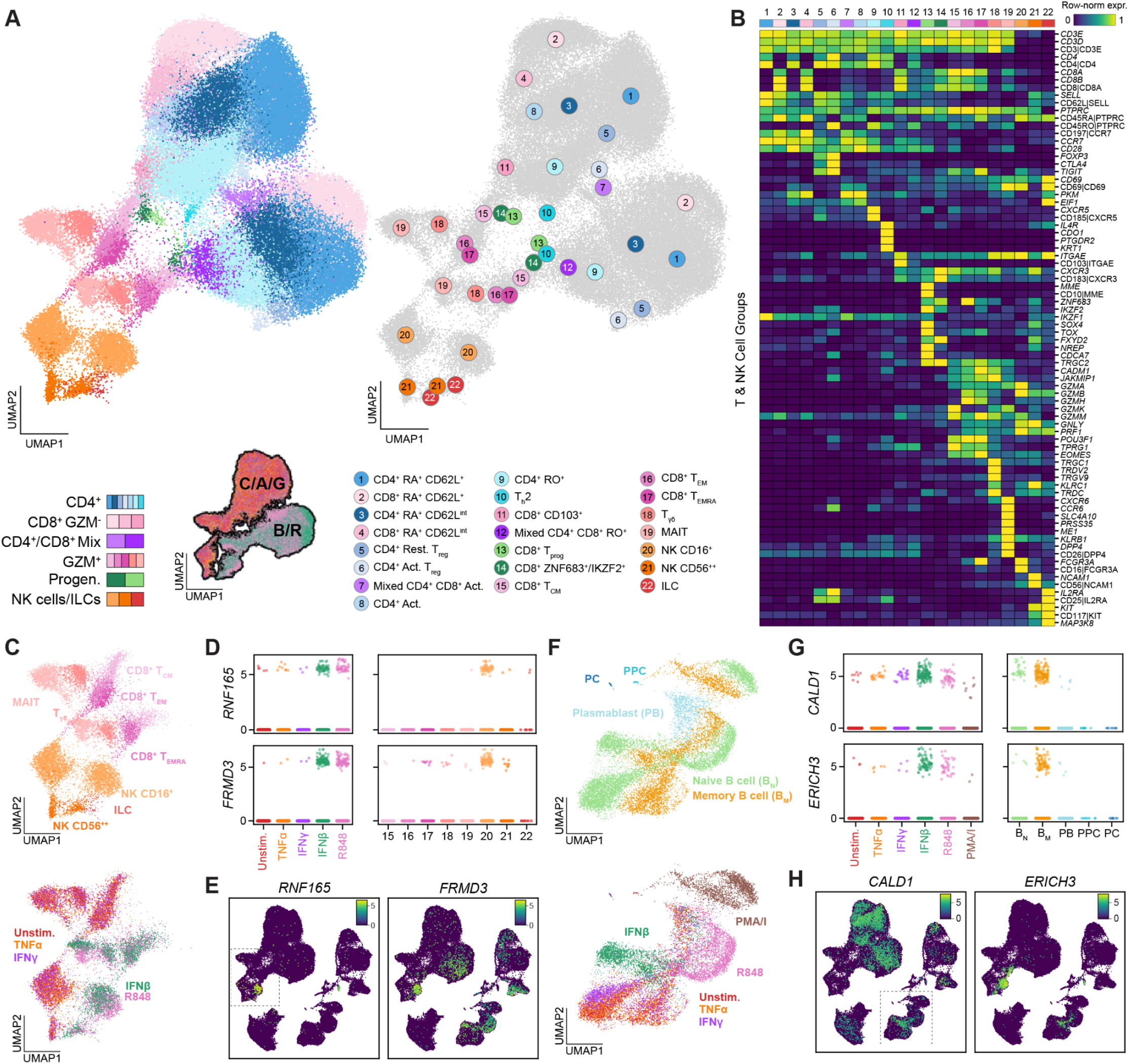
Iterative clustering and less restrictive gene filtration enable high resolution cell type and cell state map. **A**, Portion of UMAP showing T cells and NK cells, with identified cell groups colored and numbered. Insets show the location of particular cell groups and the condition overlays (C/A/G: Control, TNFɑ, IFNɣ; B/R: IFNβ, R848). **B**, Row-normalized expression heatmap of selected genes used to identify subpopulations in **A**. **C**, Portion of UMAP showing Granzyme^+^ (GZM^+^) T cell and NK cell subsets, colored by cell type (top) and condition (bottom). **D**, Expression of *RNF165* and *FRDM3*, genes expressed in both a cell type- and condition-specific manner. Plot restricted to CD16^+^ NK cells and organized by condition (left) or restricted to IFNβ stimulation and organized by cell type (right). **E**, Full single-cell UMAP showing specific expression of *RNF165* and *FRMD3*. Dashed box indicates location of GZM^+^ T and NK cells. **F**, Portion of UMAP showing B and plasma cells, colored by cell type and condition. **G**, Expression of *CALD1* and *ERICH3* as in **D**, for memory B cells by condition and IFNβ-stimulated cells by cell type. **H**, Full single-cell UMAP showing specific expression of *CALD1* and *ERICH3*.

To systematically identify cell type-specific transcriptional responses to perturbation, we ordered the DE genes by the ratio of their log_2_(FC) from control to their mean expression in all other cell types of the same condition (**Fig. S8A**, **Table S2,** Methods). For example, we identified several genes that were upregulated in IFNβ- and R848-stimulated NK cells (cluster 20) but lowly expressed in almost all other cell types (**Fig. 4C–E, Fig. S8B**). Two of the most notable genes that emerged were *RNF165* and *FRMD3*, both of which have been recently associated with worse prognosis in colorectal cancer^25, 26^ and possibly marking tumor-infiltrating NK cells.

In addition to T and NK cells, we identified 5 subtypes within the B and plasma cells, including naive and memory B cells, plasmablasts (PB), polyclonal plasmablastic cells (PPC), and mature plasma cells (PC), which were observed across all conditions (**Fig. 4F**, **Fig. S8C**). PPCs, marked by *PCNA*, *TYMS*, and *MKI67*, comprised less than 0.02% of all cells (**Fig. S8D**) and have not been described in other PBMC datasets to the best of our knowledge. This likely reflects their *in vitro* differentiation from circulating B cells in culture, consistent with previous reports of their generation from cytokine stimulation^27^. We found that PMA/I, and to a lesser extent R848, induced the expression of canonical PB genes in memory B cells (*CD226*, *MET*, *TVP23A*, *MGLL*; **Fig. S8E–F**), suggesting that these specific perturbations may be inducing early differentiation of memory B cells into PBs. Furthermore, we identified genes specifically upregulated in IFNβ-stimulated memory B cells, including the striking upregulation of *ERICH3* encoding glutamate rich protein 3, a poorly-understood vesicle-and cilium-associated gene mainly expressed in the central nervous system^28, 29^ (**Fig. 4G–H**). In addition to memory B cells, *ERICH3* was also upregulated in NK cells, CD8^+^ T memory subsets, and pDCs specifically in response to IFNβ. Outside neuronal cells, *ERICH3* has been shown to be upregulated in B cell aggregates in the meninges of the experimental autoimmune encephalomyelitis (EAE) mouse model of multiple sclerosis^30^, a disease commonly controlled with IFNβ treatment that requires B cells for efficacy^31^.

### Type I and II interferons elicit shared and specific transcriptional responses in monocytes

We next performed a focused analysis to characterize the specific and shared transcriptional responses of classical (cM) and non-classical (ncM) monocytes to type I (IFNβ) and type II (IFNγ) interferons. In response to either IFN, hundreds of genes were upregulated to similar levels in both cMs (452) and ncMs (205), including *CXCL10* and *GBP4* (log_2_(FC) > 0.5, p_adj_ < 0.05; **Fig. 5A**). We also observed genes that were more highly induced in response to IFNγ (cM: 587, ncM: 140) including *CXCL9*, IFNβ (cM: 903, ncM: 315) including *CCL8*, or exhibited opposing effects in response to the two IFNs, such as *LRRK2* and *CCL7*.

**Figure 5.**
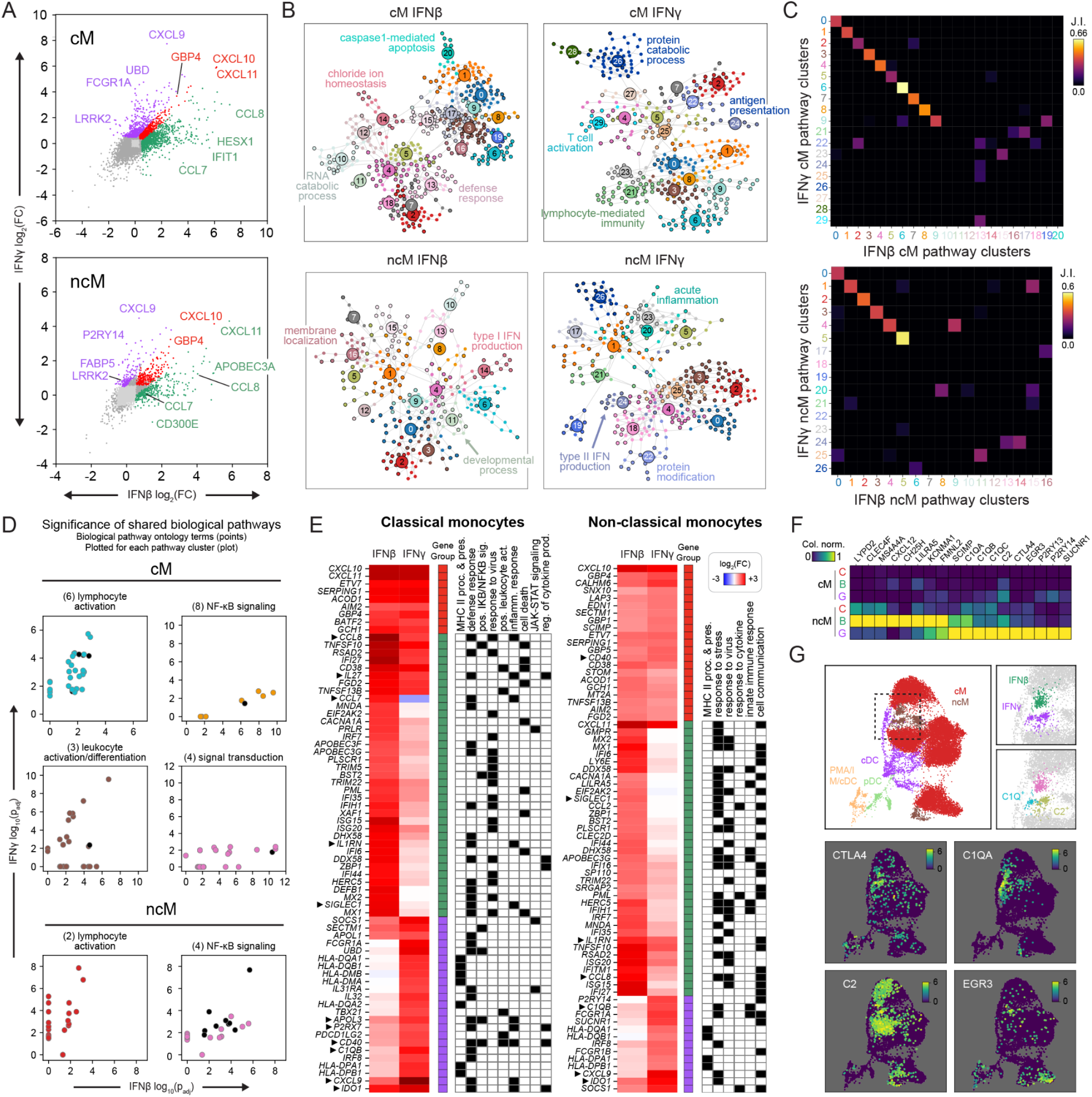
IFNs induce shared and specific transcriptional effects in classical monocytes. **A**, log_2_(FC) of gene expression from control for each IFN in classical (cM) and non-classical (ncM) monocytes. Each gene is colored by its direction of change (shared upregulated, red; IFNγ upregulated, purple; IFNβ upregulated, green). **B**, Graph of biological pathways enriched from upregulated genes for each cell type and IFN condition as determined by BiNGO. Each node is a gene ontology-enriched biological pathway term, and edges indicate shared enriched genes. Nodes are organized into “pathway clusters” via Leiden clustering using the adjacency matrix of shared genes. **C**, Jaccard index of terms between pathway clusters demonstrating some clusters are similar between the IFNs, and others are specific to either IFN. **D**, Significance (-log_10_(p_adj_)) of enriched terms comprising various shared pathway clusters in cMs (top 4 plots) and ncMs (bottom 2 plots). Unenriched terms in a given IFN have a significance set to 0. Terms are colored by their pathway cluster (title of each plot) as shown in B – C, unless they clustered differently between the IFNs, in which case they are colored black. **E**, Heatmap of log_2_(FC) for the most differentially expressed genes, organized according to direction of change as shown in **A**. Genes specific to either IFN enriched in various ontology terms are annotated with a binary matrix. **F–G**, Column-normalized heatmap and portions of UMAP showing expression of genes upregulated in IFN-stimulated non-classical monocytes.

To annotate the upregulated genes, we performed gene ontology (GO) biological pathway enrichment analysis using BiNGO^32^, which generates a network graph of enriched GO terms as nodes and shared genes between terms as edges (**Fig. 5B**). We grouped similar terms into “pathway clusters” using Leiden clustering and identified similar pathway clusters shared between the IFNs based on high Jaccard Index of ontology terms (**Fig. 5C**; Methods). In cMs, we identified 30 clusters, with 10 clusters (clusters 0–9) highly similar between the IFNs and 11 (IFNβ) and 9 (IFNγ) clusters specific to each IFN. Clusters specific to IFNβ-stimulated cells were enriched for defense response (13), chloride ion homeostasis (14), and RNA catabolic processes (1) while clusters specific to IFNγ-stimulated cells were enriched for antigen presentation (24), lymphocyte-mediated immunity (21), and protein catabolic processes (26). In ncMs, we observed 27 clusters, with 6 highly similar clusters shared between the IFNs (clusters 0–5) enriched for many of the same terms as in cMs (Jaccard Index: IFNβ, 0.397; IFNγ, 0.501), and 11 (IFNβ) and 10 (IFNγ) clusters specific to each interferon. Directly comparing the significance of terms enriched for each IFN, we note that even in highly similar pathway clusters, terms may be much more significant for one IFN than the other, including those related to lymphocyte activation in IFNγ and NF-κB signaling in IFNβ (**Fig. 5D**).

We further analyzed DE genes that may contribute to the enrichment of specific pathway terms for each IFN (**Fig. 5E**). While many genes involved in inflammatory response were similarly upregulated in cMs stimulated with either IFN, some genes exhibited specificity either in response to IFNβ, including *CCL8*, *IL27*, *CCL7*, *IL1RN*, and *SIGLEC1*, or IFNγ, including *APOL3*, *P2RX7*, *CD40*, *CXCL9*, and *IDO1*. In ncMs compared to cMs, many of the same genes and annotated pathways exhibited similar specific and shared responses to IFNβ and IFNγ. We next systematically searched for genes that exhibit an ncM-specific response to either interferon. Among the top ncM-specific genes induced by IFNβ were *CXCL12*, *CH25H*, *FMNL2*, *LILRA5,* and *KCNMA1*, all of which have been implicated in the polarization of ncMs to a migratory phenotype^33–36^ (**Fig. 5F**). In particular, *CH25H*, a known ISG with established antiviral function^37^, has been implicated in adipose-tissue inflammation in obesity and diabetes^38^. Among the top ncM-specific genes induced by IFNγ were *CTLA4*, *C1Q* complement genes*, C2, P2Y* receptors *P2RY13*, *P2RY14*, and the P2Y receptor-like *SUCNR1*. The P2Y paralogs have been previously described as ISGs in various disease and stimulation contexts^39, 40^. We note that the expression of *C1Q* and *C2* further distinguished two subpopulations of ncMs in response to IFNγ (**Fig. 5G**). *C1Q*-expressing ncMs have been reported in autoimmune diseases including systemic lupus erythematosus (SLE)^6^, while early growth response gene *EGR3* is known to be upregulated during differentiation of ncMs into macrophages and has also been implicated in autoimmune diseases with complement system dysfunction such as SLE^41, 42^. However, the induction of these populations specifically by IFNγ has not been previously reported to the best of our knowledge.

### *clue* enables the discovery of cell type-specific response expression quantitative trait loci

With its ability to encode orthogonal experimental information into each condition, the *clue* framework is uniquely suited for single-cell eQTL studies aimed to identify interactions between genetic variants and experimental conditions such as perturbations. To demonstrate this, we performed an eQTL analysis across 16 different cell types and 6 conditions, which yielded 158,445 significant *cis*-eQTLs (**Fig. 6A**). Naive CD4^+^ T cells had the highest number of eQTLs (52,016) likely reflecting the large number of cells comprising this group and the low transcriptional heterogeneity across individuals (**Fig. S9A**). Across all cell types, HLA locus genes, ribosomal proteins (e.g. *RPS26*, *RPL8*), and the aminopeptidase *ERAP2* were among the most significant eQTLs. Both shared (*PLEC*, *DNAJC15*) and cell type-specific eQTLs (*CTSW*, *ARHGAP24*, *CD151*) were observed, some of which only emerged in response to stimulation (*GBP7*, *IFITM3*, and *SLFN5*; **Fig. S9B–D**).

**Figure 6.**
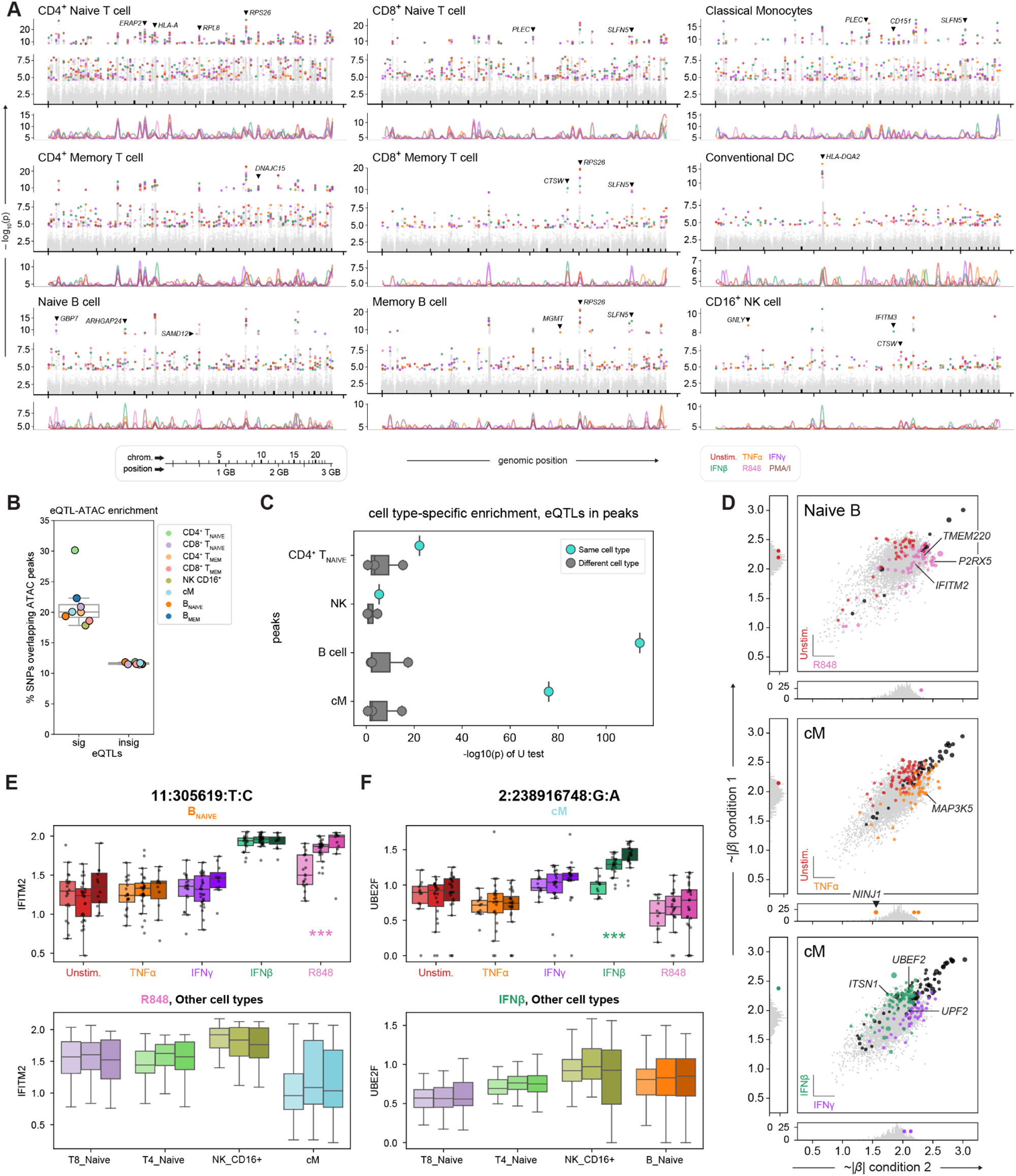
Genetic variants influence gene expression in a cell type- and condition-specific manner. **A**, Genome-wide Manhattan plots for selected cell types. All SNPs are colored gray and significant hits are colored by condition. Below each scatter plot is a line plot showing relative enrichment using a moving window average (see Methods). **B–C**, Enrichment of eQTLs in ATAC peaks, called on all unstimulated cells together (**B**) and in a cell type-specific manner (**C**, column-normalized). **D**, Comparisons of effect sizes of eQTLs between conditions in selected cell types. Significant eQTLs in either condition are colored by condition, and colored black if significant in both. SNPs that were insignificant but reported in both conditions are plotted in the main plot, colored gray. SNPs for which effect sizes were not reported in one or the other condition are plotted in the marginal distributions. **E–F**, Box plots showing eQTLs observed in a combination of cell type and condition, plotting gene expression with genotype (homozygous reference → heterozygous → homozygous alternate). Top plots show expression levels by condition in the given cell type. Bottom plots show expression levels by cell type in the given condition. Box plots showing a significant correlation (BH < 0.001) are noted with ***.

We and others have previously shown that cell type-specific *cis*-eQTLs are enriched in cell type-specific *cis-*regulatory elements. To confirm this observation, we performed enrichment analysis using cell type-specific regions of chromatin accessibility estimated from the single-cell ATAC-seq data from the AMO experiment. In unstimulated cells, *cis*-eQTLs were enriched in ATAC peaks called across all cell types (**Fig. 6B**, Methods). Furthermore, *cis*-eQTLs detected in a given cell type are significantly enriched for peaks specific to the same cell type (Mann Whitney U: CD4^+^ T_NAIVE_, *p* = 6.4 ⨉ 10^-23^; NK, *p* = 4.1 ⨉ 10^-6^; B cell, *p* = 9.7 ⨉ 10^-115^; cM, *p* = 8.3 ⨉ 10^-77^; **Fig. 6C**).

We further explored how *cis-*eQTLs could modify the effects of stimulation by comparing the effect sizes and significance for shared and condition-specific eQTLs (**Fig. 6D**). For example, we identified R848-specific *cis-*eQTLs for *TMEM220*, *IFITM2*, and *P2RX5* in naive B cells and TNFα-specific *cis-*eQTLs for *MAP3K5* and *NINJ1* in cMs. Both *MAP3K5* and *NINJ1* are known to be induced by TNFα and have been previously reported as eQTLs in lung^43^ and heart^44^. Furthermore within cMs, we observed some of the most significant *cis-*eQTLs in response to the interferons including IFNβ-specific *cis-*eQTLs for *ITSN1*, which has been previously reported in whole blood and skin, and IFNγ-specific *cis-*eQTLs for *UPF2*, a regulator of nonsense-mediated decay implicated in developmental disorders and with links to immune infiltration into the brain by macrophages and other immune cells^45^. Finally, we demonstrate that a subset of these associations are specific to both cell type and condition. For example, significant associations in *IFITM2* were found solely in R848-stimulated naïve B cells, while associations in *UBE2F* were restricted to IFNβ-stimulated cMs (**Fig. 6E–F**). These findings demonstrate the power of utilizing the *clue* framework for population-scale single-cell eQTL analyses, mapping genetic variants that interact with experimental perturbations to impact gene expression across multiple cell types.

## Discussion

Multiplexed single-cell sequencing (mux-seq) is emerging as a systematic approach to characterize the molecular profiles of cell types in large population cohorts. The integration of experimental perturbations and donor genetics enables the analysis of interindividual variability in molecular response and its genetic determinants. However, existing mux-seq implementations require reference genotyping or experimental barcoding, which incurs additional cost and may be experimentally challenging to deploy. To overcome these challenges, we developed *clue*, a framework for designing mux-seq experiments where single cells can be deterministically demultiplexed utilizing only the genotypes detected from the data. Central to *clue* is the development of *freemuxlet*, an algorithm that clusters single cells based on their genetic profiles and identifies instances where multiple cells from distinct individuals receive the same partition (droplet or well) barcode. *clue* obviates the need for reference genotyping while yielding high quality single-cell epigenomic, transcriptomic, and surface protein profiles from many individuals that can be used in studies of the genetic determinants of gene regulation.

To demonstrate the utility of the *clue* framework, we performed RNA and surface proteome sequencing in PBMCs from 64 individuals, introduced perturbations by taking advantage of redundant samples (creating 384 unique individual-conditions profiled in 12 pools), and performed differential expression and eQTL analyses with the resulting data. Genetic clustering using *freemuxlet,* followed by demultiplexing, assigned cells to individuals with high signal-to-noise and was robust to technical errors. The resulting demultiplexed data showed enrichment of differentially expressed genes and proteins in relevant biological pathways across 12 broad cell types and 6 conditions. Stimulation induced cell type and stimulation-specific expression of genes participating in inflammation, cytokine signaling, and adaptive and innate immune responses.

The analysis of our data identified rare cell types and states previously not described from scRNA-seq of PBMCs that likely developed in culture or in response to stimulation. For example, we observed several tissue-resident phenotypes in multiple CD8^+^ T cell subsets, distinguished most notably by the expression of CD103 (*ITGAE*) and *ZNF683* (which encodes HOBIT). While circulating CD103^+^ CD4+ T cells have been described in healthy individuals and proposed to be the basal recirculation of a skin-resident population^46^, their CD8^+^ counterparts have not been previously described or characterized.

We found profound cell type-specific responses to TLR and IFNAR stimulation across monocyte and lymphocyte subsets. In particular IFNβ, and to a lesser extent R848, induced high expression of *RNF165* and *ERICH3* in lymphocyte but not monocyte subtypes, genes that have been implicated in colorectal cancer and autoimmunity. IFNβ and IFNγ induced condition-specific and cell type-specific responses in classical and non-classical monocytes. Specific to non-classical monocytes, we observed that IFNβ induced a gene program suggestive of a migratory phenotype while IFNγ stimulation produced two subpopulations differentiated by the expression of complement components and *EGR3*. The two populations may correspond to recently-described subsets of ncMs distinguished by 6-sulfo LacNAc (SLAN, a carbohydrate modification of PSGL-1 protein, encoded by *SELPLG*), CD9, and CD61 surface expression^47^. We see higher albeit not statistically significant mean expression of CD9 transcript and protein, CD61 protein, and *SELPLG* transcript in the *C2*-expressing cluster, consistent with their annotations. However, further functional studies of these cell types to determine what role, if any, these genes play in the response to these agonists.

Lastly, we demonstrate the *clue* framework can be deployed for the mapping of eQTLs, demonstrate eQTL enrichment in ATAC peaks separately generated using *clue*, and explore those eQTLs that emerge only in certain cell types and stimulation conditions. We propose novel cell type- and condition-specific eQTLs in myeloid cells and B cells. We demonstrated *clue* at scale using CITE-seq but anticipate that *clue* can also be deployed for ATAC-seq and multiomic profiling of chromatin state and gene expression. While we report eQTLs identified by the integrated analysis of genotyping data, we anticipate that full-length cDNA sequencing and single-cell ATAC-seq may capture sufficient numbers of SNPs to enable high quality imputation and genetic mapping studies from single-cell genomic data alone. Indeed, emerging studies have already demonstrated that genotypes detected solely from scRNA-seq reads may be sufficient for eQTL discovery^48–50^.

There are several practical considerations for deploying the *clue* framework at scale. First, the *clue* framework is not explicitly developed to identify samples utilizing genotyping data. In fact, any multiplexing scheme can benefit from *clue* if the same barcoded samples will be profiled across multiple conditions. Second, for large experiments, we advise that statistical power be assessed carefully before employing the framework. Given a total number of cells to be sequenced for an experiment, including tens or hundreds of individuals in a pooling matrix with high compression will result in fewer cells per individual, which may hinder the ability to carry out certain downstream analyses. One way to compensate for low cell numbers per sample would be to minimize or omit cross-pool variation (e.g. no stimulation conditions). Another would be to assay the same pool in multiple single-cell reactions, though this increases overall costs. Finally, committing to assaying a large number of samples in one experiment involves some assumption of risk, especially if samples are precious. Robotics are recommended, if available, to minimize human error and experiment duration. With these considerations, *clue* is a valuable framework for highly-multiplexed single-cell sequencing studies, obviates the need for reference genotypes, can be used for both RNA and ATAC profiling, and is scalable to genetic studies involving tens or hundreds of individuals.

## Methods References

Phred-scale base quality score^51^

Detecting contamination of human DNA samples^52^ ImmVar studies^53–55^

## References

1. Freedman, M. L., et al. Principles for the post-GWAS functional characterization of cancer risk loci. Nat. Genet. 43, 513–518 (2011).

2. Blattler, A., et al. Global loss of DNA methylation uncovers intronic enhancers in genes showing expression changes. Genome Biol. 15, 469 (2014).

3. Varshney, A., et al. Genetic regulatory signatures underlying islet gene expression and type 2 diabetes. Proc. Natl. Acad. Sci. U. S. A. 114, 2301–2306 (2017).

4. Hulur, I., et al. Enrichment of inflammatory bowel disease and colorectal cancer risk variants in colon expression quantitative trait loci. BMC Genomics 16, 138 (2015).

5. Finucane, H. K., et al. Heritability enrichment of specifically expressed genes identifies disease-relevant tissues and cell types. Nat. Genet. 50, 621–629 (2018).

6. Perez, R. K., et al. Single-cell RNA-seq reveals cell type-specific molecular and genetic associations to lupus. Science 376, eabf1970 (2022).

7. Nathan, A., et al. Multimodally profiling memory T cells from a tuberculosis cohort identifies cell state associations with demographics, environment and disease. Nat. Immunol. 22, 781–793 (2021).

8. Yazar, S., et al. Single-cell eQTL mapping identifies cell type-specific genetic control of autoimmune disease. Science 376, eabf3041 (2022).

9. Fujita, M., et al. Cell-subtype specific effects of genetic variation in the aging and Alzheimer cortex. bioRxiv 2022.11.07.515446 (2022) doi:10.1101/2022.11.07.515446.

10. van der Wijst, M. G. P., et al. Single-cell RNA sequencing identifies celltype-specific cis-eQTLs and co-expression QTLs. Nat. Genet. 50, 493–497 (2018).

11. Kang, H. M., et al. Multiplexed droplet single-cell RNA-sequencing using natural genetic variation. Nat. Biotechnol. 36, 89–94 (2018).

12. Prabhu, S. & Pe’er, I. Overlapping pools for high-throughput targeted resequencing. Genome Res. 19, 1254–1261 (2009).

13. Xu, J., et al. Genotype-free demultiplexing of pooled single-cell RNA-seq. Genome Biol. 20, 290 (2019).

14. Huang, Y., McCarthy, D. J. & Stegle, O. Vireo: Bayesian demultiplexing of pooled single-cell RNA-seq data without genotype reference. Genome Biol. 20, 273 (2019).

15. Heaton, H., et al. Souporcell: robust clustering of single-cell RNA-seq data by genotype without reference genotypes. Nat. Methods 17, 615–620 (2020).

16. Weber, L. M., et al. Genetic demultiplexing of pooled single-cell RNA-sequencing samples in cancer facilitates effective experimental design. Gigascience 10, (2021).

17. Vivier, E., et al. Innate Lymphoid Cells: 10 Years On. Cell 174, 1054–1066 (2018).

18. Carvelli, J., et al. Imbalance of Circulating Innate Lymphoid Cell Subpopulations in Patients With Septic Shock. Front. Immunol. 10, 2179 (2019).

19. Cook, J. R., Craig, F. E. & Swerdlow, S. H. Benign CD10-positive T cells in reactive lymphoid proliferations and B-cell lymphomas. Mod. Pathol. 16, 879–885 (2003).

20. Hystad, M. E., et al. Characterization of early stages of human B cell development by gene expression profiling. The Journal of Immunology 182, 5882–5882 (2009).

21. Schilham, M. W., Moerer, P., Cumano, A. & Clevers, H. C. Sox-4 facilitates thymocyte differentiation. Eur. J. Immunol. 27, 1292–1295 (1997).

22. Lee, M. S., Hanspers, K., Barker, C. S., Korn, A. P. & McCune, J. M. Gene expression profiles during human CD4+ T cell differentiation. Int. Immunol. 16, 1109–1124 (2004).

23. Schattgen, S. A., et al. Integrating T cell receptor sequences and transcriptional profiles by clonotype neighbor graph analysis (CoNGA). Nat. Biotechnol. 40, 54–63 (2022).

24. Parga-Vidal L. et al. Hobit identifies tissue-resident memory T cell precursors that are regulated by Eomes. Sci Immunol 6, (2021).

25. Shembrey, C. et al. A new highly-specific Natural Killer cell-specific gene signature predicting recurrence in colorectal cancer patients. bioRxiv 2022.04.29.489868 (2022) doi:10.1101/2022.04.29.489868.

26. Chen, T.-J., et al. High FRMD3 expression is prognostic for worse survival in rectal cancer patients treated with CCRT. Int. J. Clin. Oncol. 26, 1689–1697 (2021).

27. Tarte, K., Zhan, F., De Vos, J., Klein, B. & Shaughnessy, J., Jr. Gene expression profiling of plasma cells and plasmablasts: toward a better understanding of the late stages of B-cell differentiation. Blood 102, 592–600 (2003).

28. Alsolami, M., Kuhns, S., Alsulami, M. & Blacque, O. E. ERICH3 in Primary Cilia Regulates Cilium Formation and the Localisations of Ciliary Transport and Sonic Hedgehog Signaling Proteins. Sci. Rep. 9, 16519 (2019).

29. Liu, D., et al. ERICH3: vesicular association and antidepressant treatment response. Mol. Psychiatry 26, 2415–2428 (2021).

30. Schropp, V., et al. Contribution of LTi and TH17 cells to B cell aggregate formation in the central nervous system in a mouse model of multiple sclerosis. J. Neuroinflammation 16, 111 (2019).

31. Schubert, R. D., et al. IFN-β treatment requires B cells for efficacy in neuroautoimmunity. J. Immunol. 194, 2110–2116 (2015).

32. Maere, S., Heymans, K. & Kuiper, M. BiNGO: a Cytoscape plugin to assess overrepresentation of gene ontology categories in biological networks. Bioinformatics 21, 3448–3449 (2005).

33. Sanyal, R., et al. MS4A4A: a novel cell surface marker for M2 macrophages and plasma cells. Immunol. Cell Biol. 95, 611–619 (2017).

34. Sánchez-Martín, L., et al. The chemokine CXCL12 regulates monocyte-macrophage differentiation and RUNX3 expression. Blood 117, 88–97 (2011).

35. Cheng, S.-M., et al. Differential expression of distinct surface markers in early endothelial progenitor cells and monocyte-derived macrophages. Gene Expr. 16, 15–24 (2013).

36. Lee, P. Y., et al. Type I interferon modulates monocyte recruitment and maturation in chronic inflammation. Am. J. Pathol. 175, 2023–2033 (2009).

37. Song, H., et al. Hepatitis B Virus-Induced Imbalance of Inflammatory and Antiviral Signaling by Differential Phosphorylation of STAT1 in Human Monocytes. J. Immunol. 202, 2266– 2275 (2019).

38. Russo, L., et al. Cholesterol 25-hydroxylase (CH25H) as a promoter of adipose tissue inflammation in obesity and diabetes. Mol Metab 39, 100983 (2020).

39. Zhang, C., et al. IFN-stimulated P2Y13 protects mice from viral infection by suppressing the cAMP/EPAC1 signaling pathway. J. Mol. Cell Biol. 11, 395–407 (2019).

40. Bhakta, N. R. et al. IFN-stimulated Gene Expression, Type 2 Inflammation, and Endoplasmic Reticulum Stress in Asthma. Am. J. Respir. Crit. Care Med. 197, 313–324 (2018).

41. Morita, K., et al. Emerging roles of Egr2 and Egr3 in the control of systemic autoimmunity. Rheumatology 55, ii76–ii81 (2016).

42. Carter, J. H. & Tourtellotte, W. G. Early growth response transcriptional regulators are dispensable for macrophage differentiation. J. Immunol. 178, 3038–3047 (2007).

43. Hildebrandt, M. A. T., et al. Genetic variation in the TNF/TRAF2/ASK1/p38 kinase signaling pathway as markers for postoperative pulmonary complications in lung cancer patients. Sci. Rep. 5, 12068 (2015).

44. Toma, L., et al. Ninjurin-1 upregulated by TNFα receptor 1 stimulates monocyte adhesion to human TNFα-activated endothelial cells; benefic effects of amlodipine. Life Sci. 249, 117518 (2020).

45. Johnson, J. L., et al. Inhibition of Upf2-Dependent Nonsense-Mediated Decay Leads to Behavioral and Neurophysiological Abnormalities by Activating the Immune Response. Neuron 104, 665–679.e8 (2019).

46. Klicznik, M. M. et al. Human CD4^+^CD103^+^ cutaneous resident memory T cells are found in the circulation of healthy individuals. Sci Immunol 4, (2019).

47. Kapellos, T. S., et al. Human Monocyte Subsets and Phenotypes in Major Chronic Inflammatory Diseases. Front. Immunol. 10, 2035 (2019).

48. Ma, T., Li, H. & Zhang, X. Discovering single-cell eQTLs from scRNA-seq data only. Gene 829, 146520 (2022).

49. Deelen, P., et al. Calling genotypes from public RNA-sequencing data enables identification of genetic variants that affect gene-expression levels. Genome Med. 7, 30 (2015).

50. Schadt, E. E., Woo, S. & Hao, K. Bayesian method to predict individual SNP genotypes from gene expression data. Nat. Genet. 44, 603–608 (2012).

51. Ewing, B. & Green, P. Base-calling of automated sequencer traces using phred. II. Error probabilities. Genome Res. 8, 186–194 (1998).

52. Jun, G., et al. Detecting and estimating contamination of human DNA samples in sequencing and array-based genotype data. Am. J. Hum. Genet. 91, 839–848 (2012).

53. Raj, T., et al. Polarization of the effects of autoimmune and neurodegenerative risk alleles in leukocytes. Science 344, 519–523 (2014).

54. Lee, M. N., et al. Common genetic variants modulate pathogen-sensing responses in human dendritic cells. Science 343, 1246980 (2014).

55. Ye, C. J., et al. Intersection of population variation and autoimmunity genetics in human T cell activation. Science 345, 1254665 (2014).

