## Supplemental Figures for "Reference-free multiplexed single-cell sequencing identifies genetic modifiers of the human immune response"

Figure S1

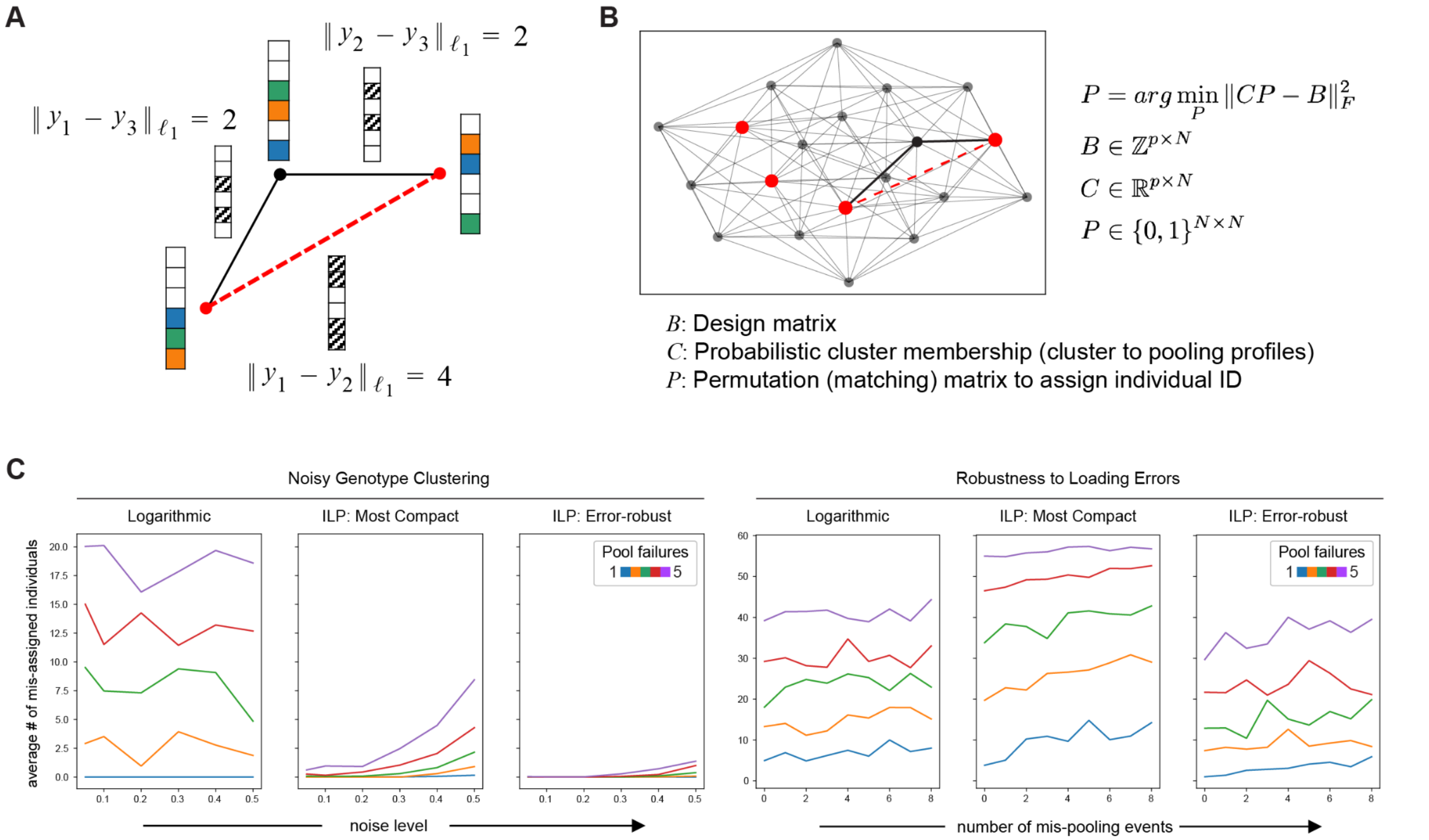


**Figure S1. Integer Linear Programming Finds the Optimal *clue* Solution. A – B**, Each donor is associated with a pooling profile, a binary vector $y$, indicating the pools into which their cells were mixed. We construct a graph with vertices representing pooling profiles, and edges connecting similar pairs of profiles within a certain Hamming distance. To tolerate errors and noise, we choose pooling profiles that are dissimilar, which correspond to anti-clique or independent sets (red vertices). Given a desired robustness, we find the maximum independent set (MIS) to achieve the most compact design. **C**, For different *clue* implementations, we demonstrate robustness to various sources of error by reporting the low average number of mis-assigned individuals given certain levels of genotype clustering noise, number of experimental mis-pooling events, and number of failures of entire pools. Genotype clustering noise is implemented as constants added randomly to the normalized probabilities of each cluster belonging to a given individual (see Methods). In general, ILP solutions are more robust to genotype clustering noise, while mis-pooling events combined with pool failures almost certainly result in some level of mis-assigned individuals regardless of implementation.

Figure S2

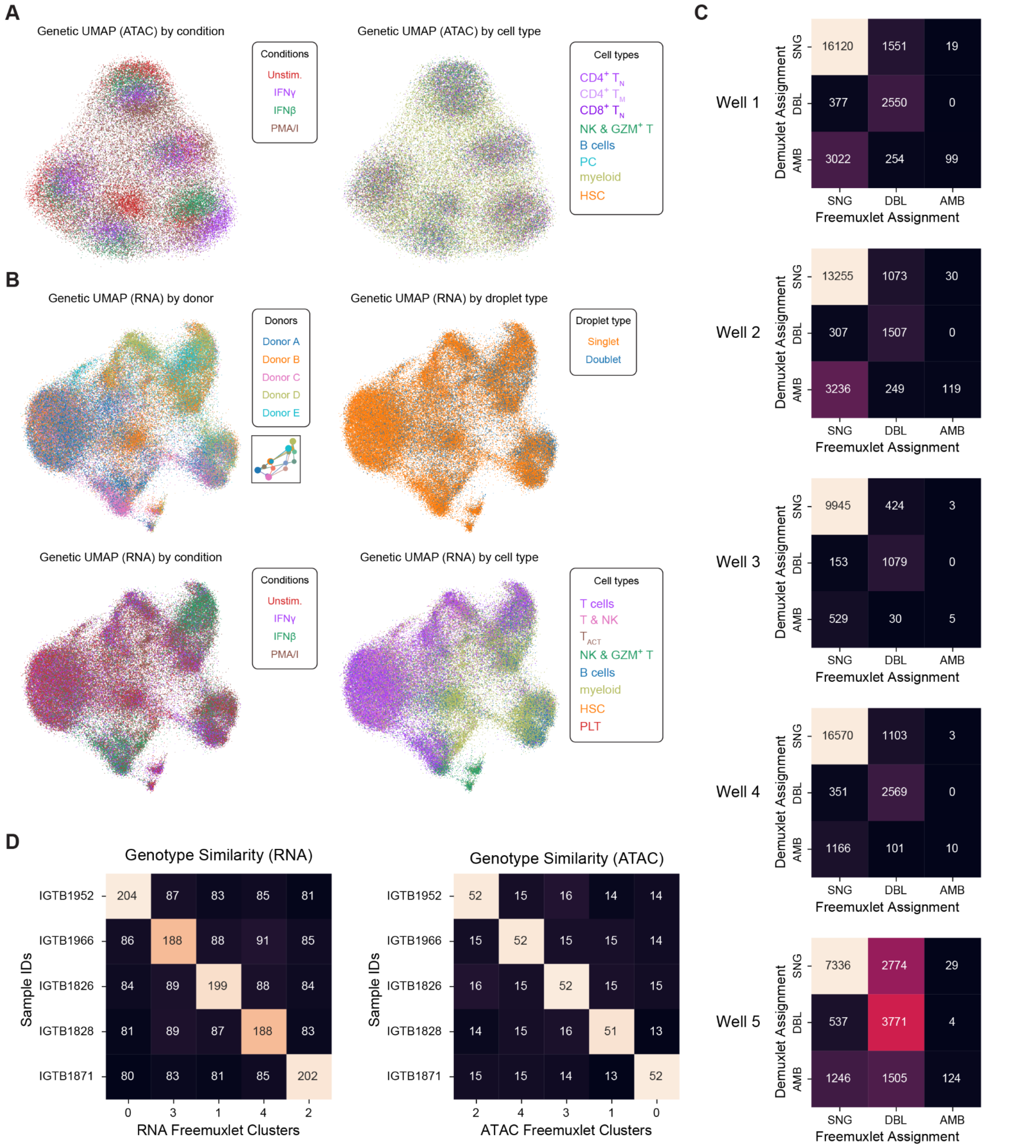


Well 1

| Demuxlet \ Freemuxlet | SNG | DBL | AMB |
|---|---|---|---|
| SNG | 16120 | 1551 | 19 |
| DBL | 377 | 2550 | 0 |
| AMB | 3022 | 254 | 99 |

Well 2

| Demuxlet \ Freemuxlet | SNG | DBL | AMB |
|---|---|---|---|
| SNG | 13255 | 1073 | 30 |
| DBL | 307 | 1507 | 0 |
| AMB | 3236 | 249 | 119 |

Well 3

| Demuxlet \ Freemuxlet | SNG | DBL | AMB |
|---|---|---|---|
| SNG | 9945 | 424 | 3 |
| DBL | 153 | 1079 | 0 |
| AMB | 529 | 30 | 5 |

Well 4

| Demuxlet \ Freemuxlet | SNG | DBL | AMB |
|---|---|---|---|
| SNG | 16570 | 1103 | 3 |
| DBL | 351 | 2569 | 0 |
| AMB | 1166 | 101 | 10 |

Well 5

| Demuxlet \ Freemuxlet | SNG | DBL | AMB |
|---|---|---|---|
| SNG | 7336 | 2774 | 29 |
| DBL | 537 | 3771 | 4 |
| AMB | 1246 | 1505 | 124 |

Genotype Similarity (RNA)

| Sample IDs | 0 | 3 | 1 | 4 | 2 |
|---|---|---|---|---|---|
| IGTB1952 | 204 | 87 | 83 | 85 | 81 |
| IGTB1966 | 86 | 188 | 88 | 91 | 85 |
| IGTB1826 | 84 | 89 | 199 | 88 | 84 |
| IGTB1828 | 81 | 89 | 87 | 188 | 83 |
| IGTB1871 | 80 | 83 | 81 | 85 | 202 |

Genotype Similarity (ATAC)

| Sample IDs | 2 | 4 | 3 | 1 | 0 |
|---|---|---|---|---|---|
| IGTB1952 | 52 | 15 | 16 | 14 | 14 |
| IGTB1966 | 15 | 52 | 15 | 15 | 14 |
| IGTB1826 | 16 | 15 | 52 | 15 | 15 |
| IGTB1828 | 14 | 15 | 16 | 51 | 13 |
| IGTB1871 | 15 | 15 | 14 | 13 | 52 |

**Figure S2. Genetic UMAPs and Concordance with Demuxlet. A**, ATAC genetic UMAP by condition and cell type. **B**, RNA genetic UMAP by donor, droplet type, condition, and cell type. **C**, Concordance of droplet assignments between demuxlet and freemuxlet, by 10x droplet reaction. **D**, Genotype similarity for RNA and ATAC.

Figure S3

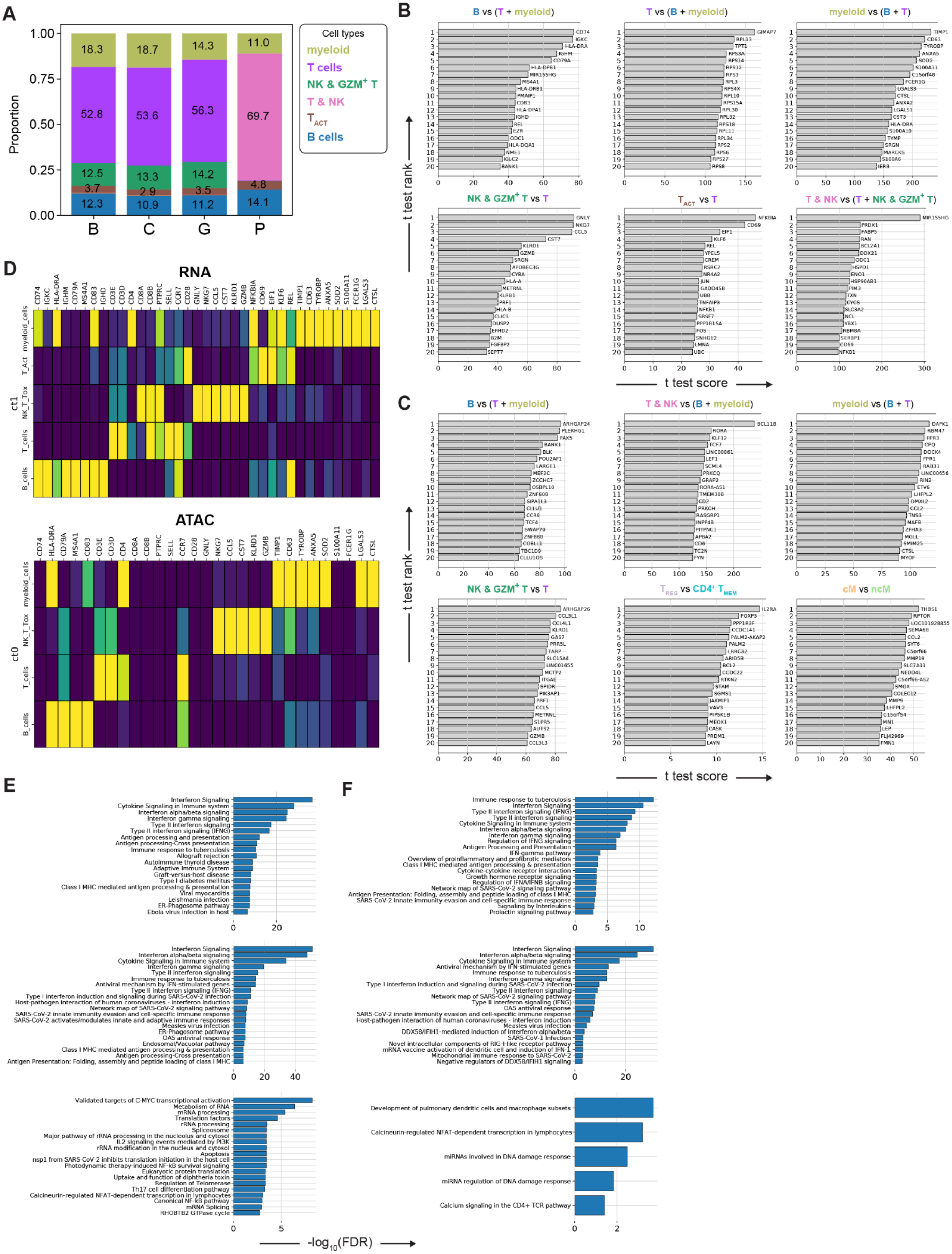
A
B
C
D
E
F
Cell types
myeloid
T cells
T & NK
B cells
Proportion
RNA
ATAC
t test rank
t test score
-log10(FDR)

**Figure S3. Differential Expression Analysis of AMO Experiment Shows Condition-specific Effects. A**, Proportion of cell types found from each condition. **B – D**, Gene rank plots (**B – C**) and heatmaps (**D**) in control conditions with top genes justifying cell type annotations, for both RNA and ATAC. Genes are ranked by Z-score underlying the BH-corrected *p* values from a *t* test of the compared cell type groups. **E – F**, Gene set enrichment results for the top 100 DE genes (using method above, comparing condition to control across all cell types). Genes shown for RNA (**E**) and ATAC (**F**) confirm the effects from stimulation conditions (top, IFNɣ; middle, IFNβ; bottom, PMA/I).

Figure S4

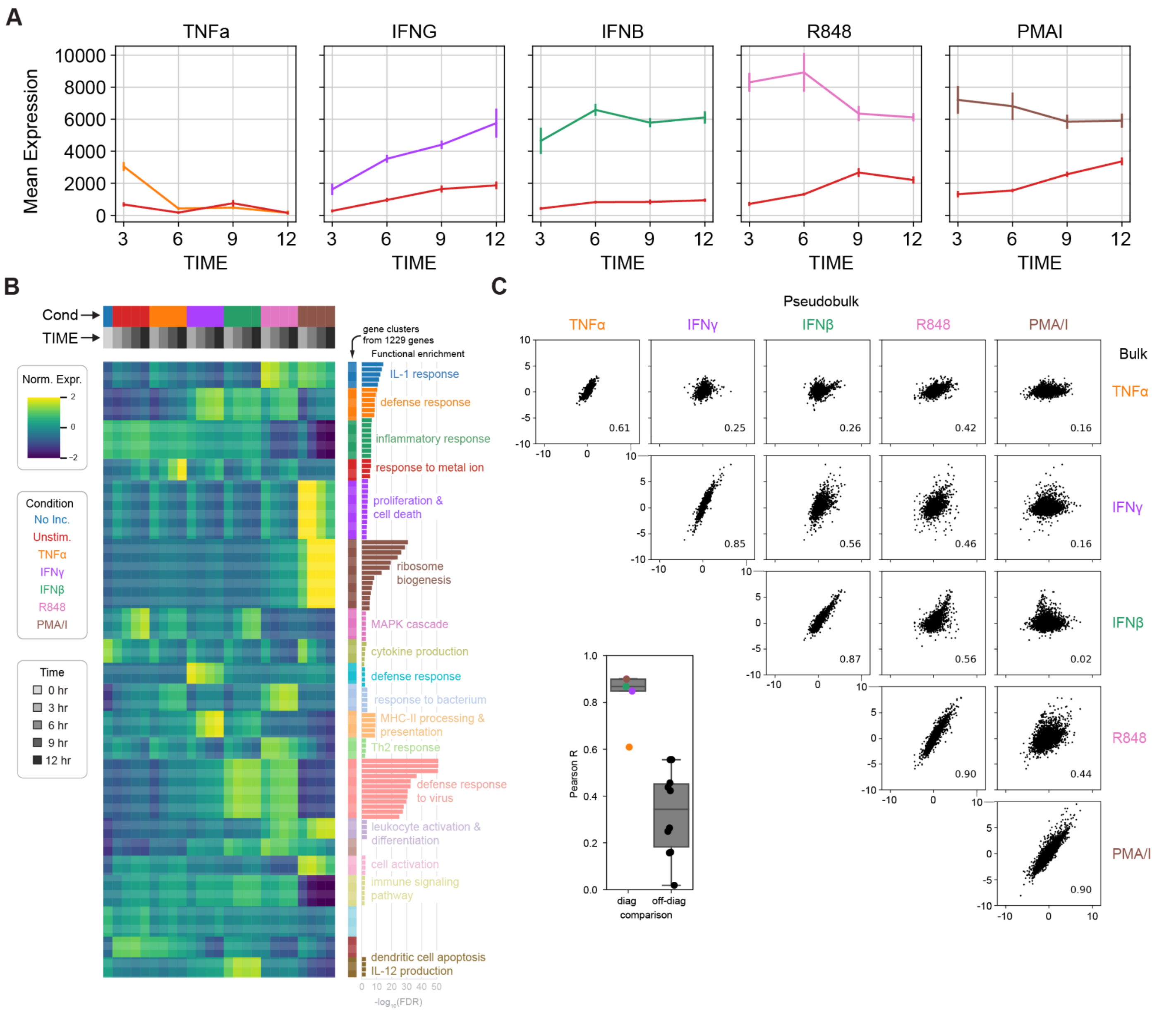


**Figure S4. Bulk RNA Sequencing Justifies Timepoints, Concordant with Single-cell Pseudobulks. A**, Line plots showing mean expression of DE genes, at each timepoint, for 4 timepoints. **B**, Heatmap of differentially expressed genes comparing all timepoints to the 0 time point for all conditions. Genes (1229) are clustered into modules along the y axis, and significance of enriched gene ontology terms plotted with bars, with top enriched term shown. **C**, Pair plot showing the correlation of normalized expression between conditions in the bulk experiment and pseudobulked profiles from the single-cell experiment. Inset: correlations for the same conditions were higher than those for different conditions, with TNF-α lower than the rest, likely due to the change in agonist concentration between the bulk and RNA-seq experiments.

Figure S5

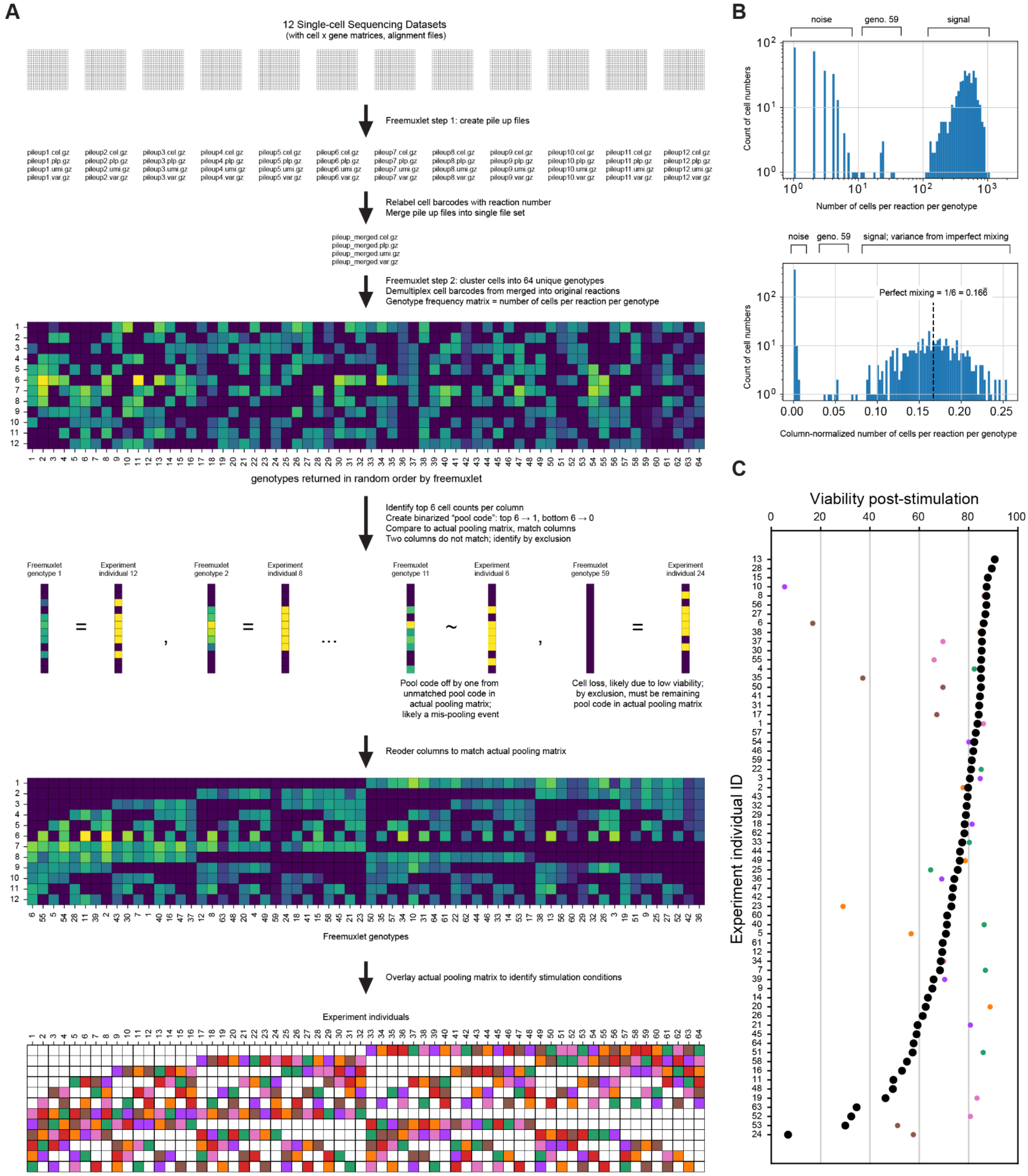
A
12 Single-cell Sequencing Datasets
(with cell x gene matrices, alignment files)
Freemuxlet step 1: create pile up files
pileup1.cel.gz
pileup1.plp.gz
pileup1.umi.gz
pileup1.var.gz
Relabel cell barcodes with reaction number
Merge pile up files into single file set
pileup_merged.cel.gz
pileup_merged.plp.gz
pileup_merged.umi.gz
pileup_merged.var.gz
Freemuxlet step 2: cluster cells into 64 unique genotypes
Demultiplex cell barcodes from merged into original reactions
Genotype frequency matrix = number of cells per reaction per genotype
genotypes returned in random order by freemuxlet
Identify top 6 cell counts per column
Create binarized "pool code": top 6 → 1, bottom 6 → 0
Compare to actual pooling matrix, match columns
Two columns do not match; identify by exclusion
Freemuxlet genotype 1
Experiment individual 12
Freemuxlet genotype 2
Experiment individual 8
Freemuxlet genotype 11
Experiment individual 6
Freemuxlet genotype 59
Experiment individual 24
Pool code off by one from unmatched pool code in actual pooling matrix; likely a mis-pooling event
Cell loss, likely due to low viability; by exclusion, must be remaining pool code in actual pooling matrix
Reoder columns to match actual pooling matrix
Freemuxlet genotypes
Overlay actual pooling matrix to identify stimulation conditions
Experiment individuals
B
noise
geno. 59
signal
Count of cell numbers
Number of cells per reaction per genotype
signal; variance from imperfect mixing
Perfect mixing = 1/6 = 0.166
Column-normalized number of cells per reaction per genotype
C
Viability post-stimulation
Experiment individual ID

**Figure S5. Deconvolution of Pooling Matrix and Viability Assessment. A**, Deconvolution of the pooling matrix to match freemuxlet genotypes to experimental individual IDs. Justifications for assigning the mispooled sample and the sample that had significant cell loss. **B**, Distribution of cell number per element in the output pooling matrix (top, absolute; bottom, column-normalized) **C**, Viability post-stimulation of selected samples as measured using flow cytometry propidium iodide staining. Black dots are the control (unstimulated) sample for every individual, while a random cross-section of other conditions were taken from a subset of individuals. Experiment ID 24's control sample, which showed significant cell loss during genotype clustering, had the lowest viability of all samples.

### Figure S6

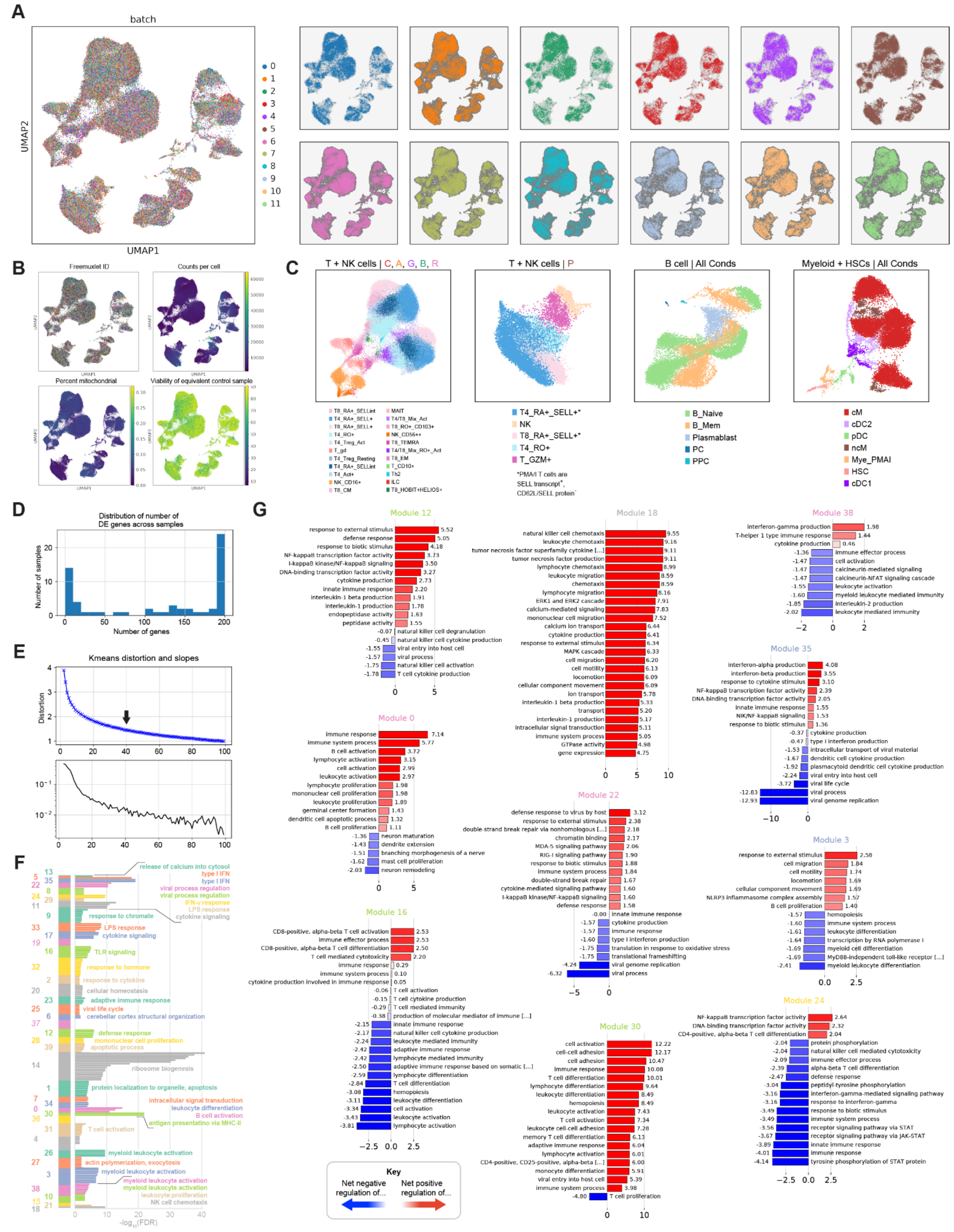

A
batch
UMAP2
UMAP1
0
1
2
3
4
5
6
7
8
9
10
11
B
Freemuxlet ID
Counts per cell
Percent mitochondrial
Viability of equivalent control sample
C
T + NK cells | C, A, G, B, R
T + NK cells | P
B cell | All Conds
Myeloid + HSCs | All Conds
T8_RA+_SELLint
T4_RA+_SELL+
T8_RA+_SELL+
T4_RO+
T4_Treg_Act
T_gd
T4_Treg_Resting
T4_RA+_SELLint
T4_Act+
NK_CD16+
T8_CM
MAIT
T4/T8_Mix_Act
T8_RO+_CD103+
NK_CD56++
T8_TEMRA
T4/T8_Mix_RO+_Act
T8_EM
T_CD10+
Th2
ILC
T8_HOBIT+HELIOS+
T4_RA+_SELL+*
NK
T8_RA+_SELL+*
T4_RO+
T_GZM+
*PMA/I T cells are SELL transcript+, CD62L/SELL protein-
B_Naive
B_Mem
Plasmablast
PC
PPC
cM
cDC2
pDC
ncM
Mye_PMAI
HSC
cDC1
D
Distribution of number of DE genes across samples
Number of samples
Number of genes
E
Kmeans distortion and slopes
Distortion
F
release of calcium into cytosol
type I IFN
type I IFN
viral process regulation
viral process regulation
IFN-γ response
LPS response
cytokine signaling
response to chromatin
LPS response
cytokine signaling
TLR signaling
response to hormone
response to cytokine
cellular homeostasis
adaptive immune response
viral life cycle
cerebellar cortex structural organization
defense response
mononuclear cell proliferation
apoptotic process
ribosome biogenesis
protein localization to organelle, apoptosis
intracellular signal transduction
leukocyte differentiation
B cell activation
antigen presentation via MHC-II
T cell activation
myeloid leukocyte activation
actin polymerization, exocytosis
myeloid leukocyte activation
myeloid leukocyte activation
myeloid leukocyte activation
leukocyte proliferation
NK cell chemotaxis
-log10(FDR)
G
Module 12
Module 18
Module 38
Module 35
Module 0
Module 22
Module 3
Module 16
Module 30
Module 24
Key
Net negative regulation of...
Net positive regulation of...

**Figure S6. The *clue_logarthmic* Framework Yields High-quality Single-cell Profiles from Stimulation Conditions. A**, UMAP colored by batch (12 droplet reactions), showing minimal technical effects. **B**, UMAP colored by freemuxlet ID, counts per cell, percent mitochondrial content, and viability of the equivalent control sample (per individual), showing minimal effects due to these covariates. **C**, Zoomed-in sections of UMAP space showing granular cell type annotations. **D**, Distribution of number of DE genes across samples (cell type + condition) included in the heatmap in Fig 3. A limit of 200 genes per sample was imposed. **E – F**, K-means sum of square errors (distortion), and slopes between adjacent points, for heatmap in Fig 3. A value of 40 was chosen where slopes become more noisy, and we empirically see (**F**) the smaller clusters formed are significant for gene ontology (GO) terms. **G**, For only those GO terms prefixed with “negative/positive regulation of”, significance values are added to estimate a “net effect” of regulation.

Figure S7

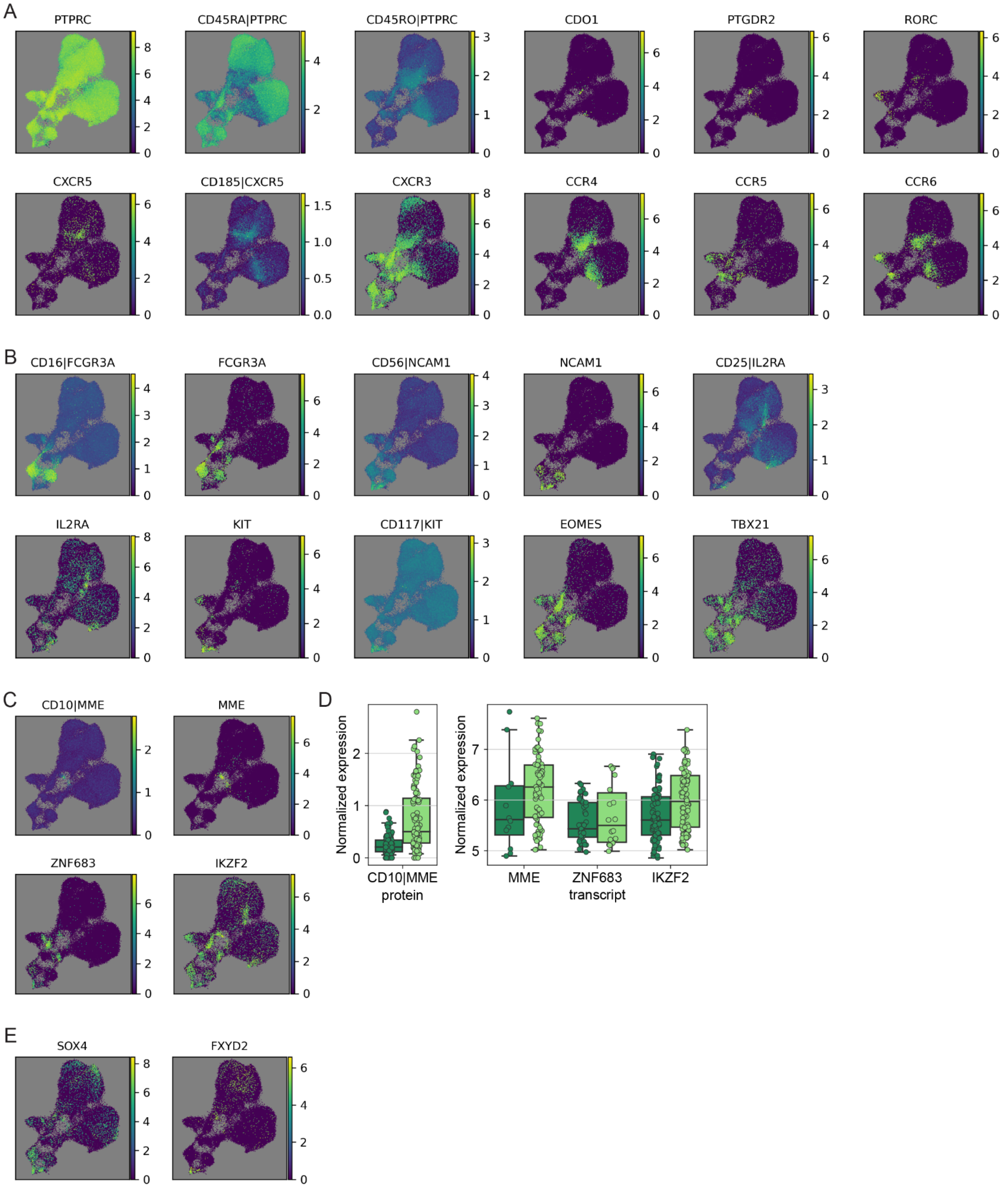
A
PTPRC
CD45RA|PTPRC
CD45RO|PTPRC
CDO1
PTGDR2
RORC
CXCR5
CD185|CXCR5
CXCR3
CCR4
CCR5
CCR6
B
CD16|FCGR3A
FCGR3A
CD56|NCAM1
NCAM1
CD25|IL2RA
IL2RA
KIT
CD117|KIT
EOMES
TBX21
C
CD10|MME
MME
ZNF683
IKZF2
D
Normalized expression
CD10|MME
protein
MME
ZNF683
IKZF2
transcript
E
SOX4
FXYD2

**Figure S7. T Lymphocytes and NK Cells Show Expression of Specific Genes and Surface Proteins. A**, Genes and proteins identifying T cell subsets. $CD45RO^+$ $CD4^+$ subsets show expression of some phenotyping markers but do not cluster out separately using Leiden. **B**, Genes and proteins identifying NK cell and ILC subsets. **C – E**, Two populations of T cells are identified by high expression of CD10 and *MME*, and show expression of HOBIT (*ZNF683*) and and HELIOS (*IKZF2*), along with other markers (*SOX4*, *FXYD2*) suggesting a progenitor-like phenotype.

Figure S8

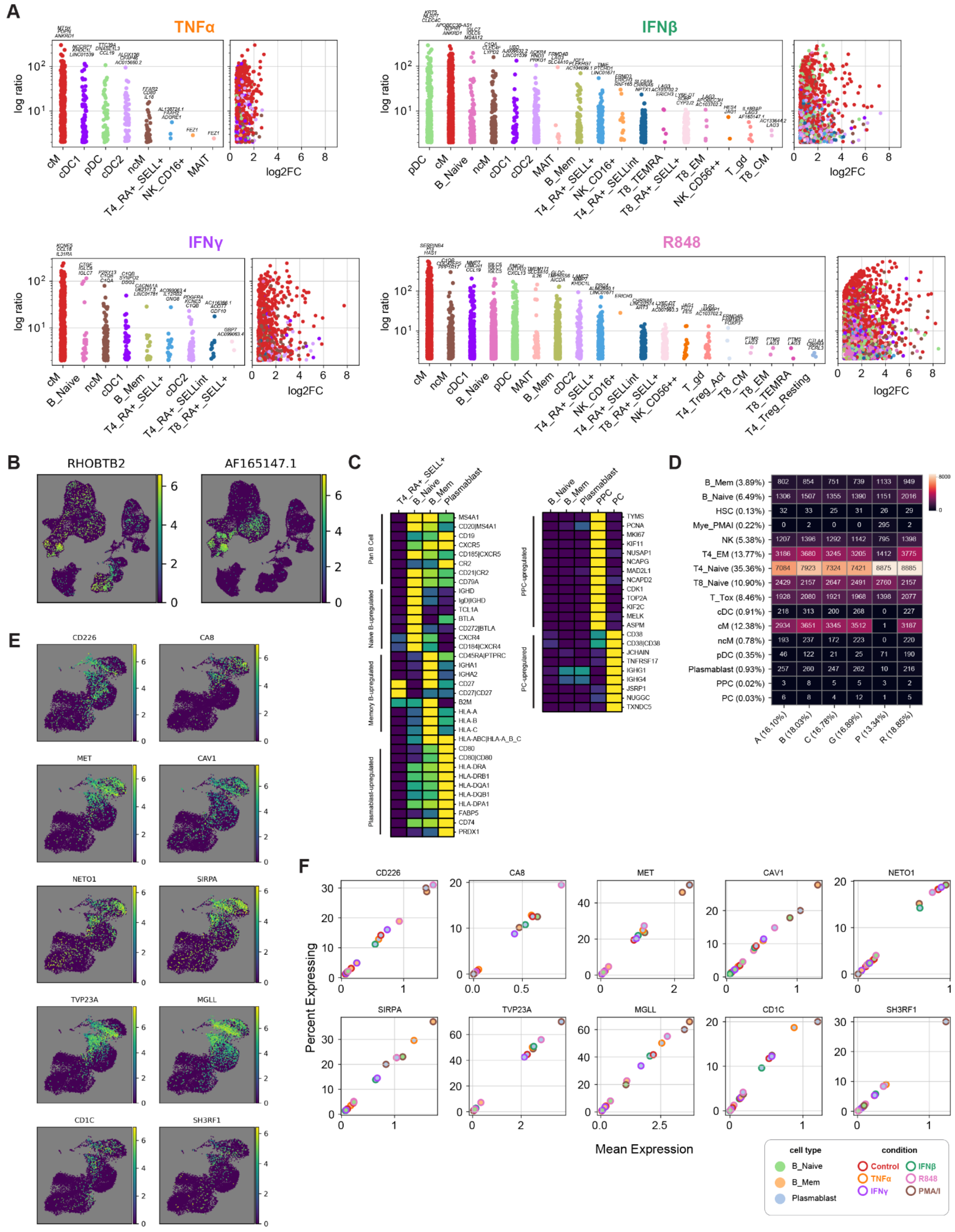
A
TNFα
IFNβ
IFNγ
R848
log ratio
log2FC
B
RHOBTB2
AF165147.1
C
Pan B Cell
Naive B-upregulated
Memory B-upregulated
Plasmablast-upregulated
PPC-upregulated
PC-upregulated
D
E
CD226
CA8
MET
CAV1
NETO1
SIRPA
TVP23A
MGLL
CD1C
SH3RF1
F
Percent Expressing
Mean Expression
cell type
B_Naive
B_Mem
Plasmablast
condition
Control
TNFα
IFNγ
IFNβ
R848
PMA/I

**Figure S8. Overview of Cell-type-specific Transcriptional Responses to Perturbation. A**, For each condition, the DE genes were ordered by the ratio of their $\log_2$(FC) from control to their mean expression in all other cell types of the same condition. This helped prioritize gene expression patterns specific to both cell type and perturbation. **B**, *RHOBTB2* and *AF165147.1*, two other genes found upregulated in a IFN-β/R848-stimulated NK cell-specific manner. **C**, Heatmap showing row-normalized gene expression used to identify B cell and Plasmablast subsets, with $CD45RA^+$ $CD4^+$ *SELL*$^+$ T cells for comparison. **D**, Absolute number and percentage of cells annotated as each cell type and condition. **E – F**, UMAPs and scatter plots showing genes upregulated in PMA/I- and R848-stimulated memory B cells, but mainly expressed only in plasmablasts of other conditions.

Figure S9

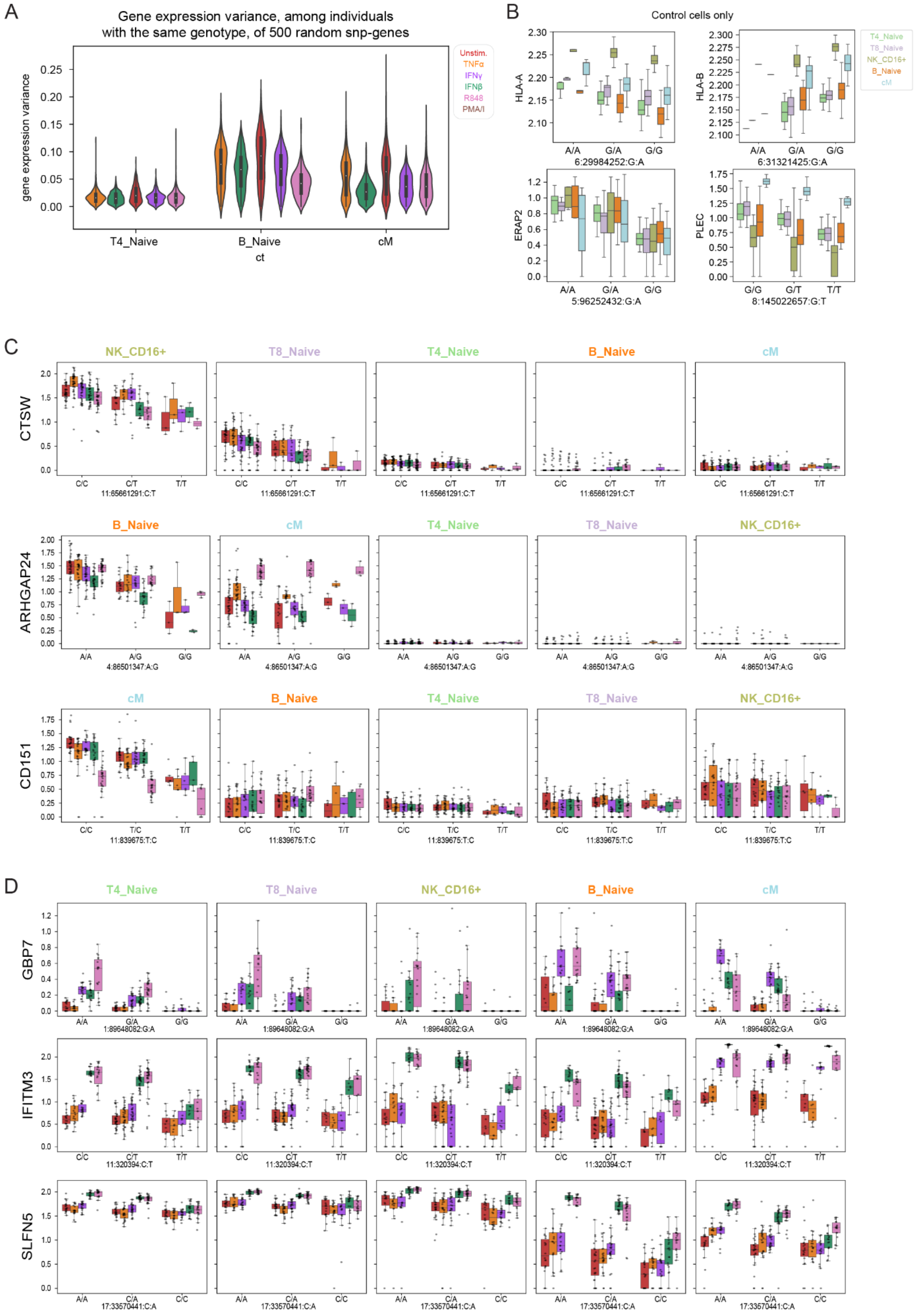
A
Gene expression variance, among individuals with the same genotype, of 500 random snp-genes
gene expression variance
Unstim.
TNFα
IFNγ
IFNβ
R848
PMA/I
T4_Naive
B_Naive
cM
ct
B
Control cells only
HLA-A
6:29984252:G:A
HLA-B
6:31321425:G:A
T4_Naive
T8_Naive
NK_CD16+
B_Naive
cM
ERAP2
5:96252432:G:A
PLEC
8:145022657:G:T
C
CTSW
11:65661291:C:T
ARHGAP24
4:86501347:A:G
CD151
11:839675:T:C
NK_CD16+
T8_Naive
T4_Naive
B_Naive
cM
D
GBP7
1:89648082:G:A
IFITM3
11:320394:C:T
SLFN5
17:33570441:C:A

**Figure S9. Overview of Cell-type- and Condition-specific eQTLs. A**, Variance in gene expression for selected cell types. For 500 random snp-gene pairs, for each cell type and condition, cells were pseudobulked by individual, then grouped by genotype and variance was calculated. Naive T cells show lower variance (transcriptional homogeneity) as compared to B cells and myeloid cells. **B**, Significant eQTLs observed across most or all cell types in control cells. **C**, Significant eQTLs emerging across most or all conditions in $CD16^{+}$ NK Cells (*CTSW*), Naive B cells (*ARHGAP24*), and cMs (*CD151*), but not in other cell types. **D**, Significant eQTLs emerging only under stimulation in most or all cell types.
