## Supplementary material for "Reference-free multiplexed single-cell sequencing identifies genetic modifiers of the human immune response": Methods

### Methods and Procedures

#### Encoding and Decoding Algorithms

While the AMO design benefits from the elimination of reaction-wise batch effect and the lack of need for independent genotyping, the design still requires as many reactions as there are individuals. Batch effect reduction can be achieved with standard multiplexing techniques, and the cost benefits of eliminating independent genotyping are quickly outweighed by the scaling of the number of reactions with the number of individuals. For that reason, we implemented the clue_logarithmic scheme, that dramatically reduces the number of reactions required to profile a given number of individuals. To construct a screen according to this code, every sample ID [0 : N-1] is converted into a bit-word - a binary representation of a sample ID number with a length $\log_2 N$, this vector is further appended by an orthogonal binary vector of the same length. Resulting vector represents an $n^{th}$ column of a sensing matrix $A^{P\times N}$, and this process is repeated for every sample of $N$. Resulting binary sensing matrix has $P = 2\log_2 N$ rows and serves as a guidance whether a sample $x_n$ has to be added into the pool $y_p$ ("1") or not ("0"). The design, based on logarithmic encoding (Algorithm 1), allows for the multiplexing of $N$ individuals into $2\log_2 N$ pools and the profiling up to $\log_2 N$ conditions per individual. Moreover, the design preserves the ability to simultaneous genotyping and profile single-cells, as well as map anonymous donors back to extra-experimental phenotypes. We further aim to assign $C = \log_2 N$ conditions and distribute them throughout the pooling scheme to maximize variance of conditions within a pool and minimize variance of condition compositions between pools. First we approach this problem by a brute force search: Algorithm 2. We set a variance threshold that represents variance between numbers of occurrences of every condition in a pool. For example, ideally if every condition is represented equally, variance will be zero, but in most cases this is very hard to achieve. So we set a default to be 1.6

Decoding of a compressed screen means definitive association of a positive read out with the sample $n$. Decodability of a linear code such as the logarithmic encoding used here is defined by disjunctness of a constructed sensing matrix $A^{P\times N}$. A sensing matrix $A^{P\times N}$ is $d$-disjunct if for any subset of columns $H \subset [0 : N-1]$ s.t. $|H| = d$ and any column $n \in [0 : N-1] - H$ there exists a row $p$ s.t. $p^{th}$ column has "1" and all columns of $H$ have "0" in columns $p$. Here $|H|$ means cardinality of a subset $H$. Logarithmic encoding produces a $d$-disjunct matrix with $d = 1$, meaning it allows definitive decoding only if the sparsity $S$ of an input is 1 which is satisfied by our experimental set up (every *freemuxlet* cluster is associated with only one genotype). In the condition when $d = S$ a simple pattern consistency decoder (PCD) can be used. PCD can be formulated as such: *for every freemuxlet cluster $\phi_i$ find sample n which corresponds to max of:*

$$\sum_{p=0}^{P-1} A_{pn} y_p$$

, where $y_p \in \{0,1\}^P$ is whether a cluster $\phi_i$ is detected in the pool $p$.

---

**Algorithm 1:** : encoding with logarithmic signatures

---

**Result:** Pooling matrix $A^{P\times N}$
**for** *every individual* $x_n \in [0 : N-1]$ **do**
  vectorN = convert $x_n$ to binary with length $\log_2 N$;
  vectorN.append(orthogonal vectorN with length $\log_2 N$);
  insert vectorN as $n^{th}$ columns of matrix $A^{P\times N}$;
**end**

---

**Algorithm 2:** : condition assignments

**Result:** Pooling binary matrix $A^{P \times N}$ with condition assignment
construct a list of permutations of conditions with length $\log_2 N$;
Set a variance threshold: thresh = 1.6;
**for** *every column p of* $A^{P \times N}$ **do**
  **while** *variance in any pool* $\geqslant$ *thresh* **do**
    **try:**
      *randomly choose a permutation;*
    **catch** *list of permutations is parsed and none satisfies "while"***:**
      *p -= 1;*
      *continue;*
    **if** $p < 0$ **then**
      *break a "while" loop and start over;*
      *p = 0;*
    **end**
    **else**
      *assign conditions using a chosen permutation to a column of* $A^{P \times N}$ *where element* $a_{pn} = 1$*;*
      *remove this permutation from the list of permutations;*
      *count current occurrences of each condition per pool;*
      *estimate variance in each pool;*
    **end**
  **end**
**end**

### Integer Linear Programming: Optimization of Compression and Error Tolerance

#### Select pooling profiles from $m$-robust candidate set

As described in the main text, we solve the maximum independent set (MIS) problem and find the largest set containing $T$ individual pooling profiles where the minimum difference between any pair in this set is $m + 1$. The MIS by construction guarantees that any size-$N$ subset can be chosen from it and assigned to the $N$ donors, and the resulting scheme can tolerate missing information in up to $m$ wells and is robust against other experimental noise. Finding the optimal subset to form the scheme is formulated into a convex optimization problem with constrains.

Consider a size-$T$ binary (selection) vector $x \in 0, 1^T$ to optimize: $x_i = 1$ if $y_i \in \{0,1\}^P$, the $i$-th pooling configuration, is included in the final scheme. Objective function to minimize:

$$\sum_{(i,j):|y_i - y_j| = m+1} x_i x_j \text{ (minimize the number of closest pairs)}$$

Satisfying:

$$\sum_{i=1}^{T} x_i = N; \quad \text{and} \quad \sum_{i=1}^{T} y_{ip} \cdot x_i \geq \lfloor \frac{NC}{P} \rfloor, \quad \forall p \in [P]$$

(Where $y_{ip}$ is the $p$-th element in $y_i$, known after the previous step.)

To linearize the quadratic objective function, we introduce $T(T-1)/2$ augmented variables, where $x_{ij} = x_i x_j \ \forall i > j$, indicating if both $y_i$ and $y_j$ are chosen. Let $\delta_{ij} = |y_i - y_j| = m + 1$, indicating if this pair of profiles is among the closest pairs. $\delta_{ij}$'s are pre-computed for all the $T(T-1)/2$ pairs. The system is then transformed into an integer (binary) linear system

$$\text{Objective: minimize } \sum_{i>j} \delta_{ij} x_{ij} \quad \textbf{Equation } 1\text{a.}$$

$$\text{Satisfying: } \sum_{i=1}^{T} x_i = N \quad \textbf{Equation } 1\text{b.}$$

$$\sum_{i=1}^{T} y_{ip} \cdot x_i \geq \lfloor \frac{NC}{P} \rfloor, \quad \forall p \in [P] \quad \textbf{Equation } 1\text{c.}$$

$$x_{ij} - x_i \leq 0; \quad x_{ij} - x_j \leq 0; \quad x_i + x_j - x_{ij} \leq 1 \quad \textbf{Equation } 1\text{d.}$$

where condition 1d is the linear equivalence to $x_{ij} = x_i x_j$ since $x_i, x_j \in 0, 1$.

#### Finding near-optimal starting point

This system can be solved by integer linear programming (ILP), but the number of parameters and therefore the searching space can be prohibitively huge (natively $O(2^{T^2/2})$), thus finding the optimal solution is hard in practice. We introduce an approach to reduce the searching space to only a fraction while achieving the optimality.

Similar as in the first step, we construct a new graph $G$ with its vertex set being the $T$ profiles. An edge exists between $y_i$ and $y_j$ if and only if they are among the closet pairs, $\|y_i - y_j\|_{\ell_1} \leq m + 1$. Those types of pairs differ by only $m + 1$ wells, the least in the candidate set, thus we want as few of them to co-exist as possible in our final scheme. Recall that if $m$-robustness is the best we can achieve with the fixed $(N, C, P)$, there is no independent set in $G$ that is larger than $N$ (otherwise we can choose that set and achieve (m+1)-robustness). Intuitively, we want to prioritize the (multiple, small) (m+1)-independent sets in our scheme, as individuals inside such a set are further apart from each other thus safer to co-exist.

We proceed as follows: find a MIS in $G$, delete it from the graph, continue to find a MIS in the remaining graph and repeat the process until we have a sequence of disjoint (m+1)-independent sets jointly covering all the $T$ vertices in original $G$. Starting from the largest (and while the total size is below $N$), we fix the top MISs to be included in the solution of the ILP by fixing the corresponding $x_i$ to 1, then optimize the system to find the rest such that we fill the $P$ wells.

#### Assigning experimental conditions to each sample

If the number of samples under each condition differs across wells, batch effect may introduce confounding in downstream analysis. A balanced scheme with approximately $\frac{N}{P}$ samples for each conditions in every well is thus preferred. Let $x_{i,p,c} \in 0, 1$ indicating if a sample from individual $i$ in condition $c$ is in well $p$, the constrain formalizes as follow, also an ILP system. Because of the inherent symmetry in the problem, we can find feasible solutions in most of the cases, and if no feasible solution is found, we relax the same-well-size constrain (2d) or the minimum batch effect constrain (2c) incrementally by the minimum step size (implemented in the software).

$$\sum_{c=1}^{C} x_{i,p,c} = y_{ip} \;\forall i,p \quad \text{(Each individual has at most one sample in each well)} \qquad \textbf{Equation } 2a.$$

$$\sum_{p=1}^{P} x_{i,p,c} = 1 \;\forall i,c \quad \text{(Each individual has exactly one sample from each condition)} \qquad \textbf{Equation } 2b.$$

$$\lfloor \frac{N}{P} \rfloor \leq \sum_{i=1}^{N} y_{ip} \cdot x_{i,p,c} \leq \lfloor \frac{N}{P} \rfloor + 1 \;\forall p,c \quad \text{(Uniform distribution of conditions over pools)} \qquad \textbf{Equation } 2c.$$

$$\sum_{i=1}^{N} \sum_{c=1}^{C} x_{i,p,c} \geq \lfloor \frac{NC}{P} \rfloor \;\forall p \quad \text{(Each well has roughly the same number of samples)} \qquad \textbf{Equation } 2d.$$

### Freemuxlet

#### Detailed Procedures of Freemuxlet

The algorithms for *freemuxlet* can be divided into two parts. The first part, described in Algorithm 3, starts with calculating the Singlet Score each droplet and reorder the droplets from largest to smallest singlet scores to ensure that multiplets do not form a cluster on its own. Next, starting from the droplets with largest singlet scores, each droplet is assigned to a new cluster, an existing cluster, or none of the clusters. If the genetic similarity between a droplet and an existing cluster is above a certain threshold of Bayes Factor $t_{BF}$, the droplet is assigned to the cluster. If no such cluster exists, the droplet forms a new cluster on its own only if the total number of existing clusters is less than the number of sample $N$. Otherwise, the droplet is not assigned to any cluster to conservatively assign only putative singlets into clusters.

The next steps, described in Algorithm 4 are very similar to the steps of *demuxlet*, with two key differences. First, instead of using genotypes obtained from external genotypes, we assume that $\Pr(g_v; C, \boldsymbol{f})$ approximate the genotype probabilities of individuals represented by each cluster $C$. Second, we perform this steps iteratively, by classifying each droplets into singlet, doublet, or ambiguous categories and updating each cluster to contain only putative singlets. The thresholds to classify droplets in our experiment is chosen consistent to demuxlet ($t_{singlet} = t_{doublet} = 2$), but can also determined by Bayes rule given specific priors of singlets and doublets. This iterative procedure is repeated for a fixed number of times (10 was used in our experiment), or until there is no change in the assignment of each droplet, whichever occurs earlier.

**Algorithm 3:** Initial clustering of droplets

**Result:** Clusters $C[1], \ldots, C[N]$ as exclusive subsets of $\{1, \ldots, D\}$
(Droplets are unordered initially.)
**for** $d \leftarrow 1$ **to** $D$ **do**
  $S[d] \leftarrow \log [L_1(d; \boldsymbol{f})] - \log [L_2(d; \alpha = 0.5, \boldsymbol{f})]$
**end**
Sort droplets in decreasing order of $S[d]$ (i.e. $S[i] \geq S[j]$ if $i < j$)
Initialize $C[1], C[2], \ldots, C[N]$ as empty sets.
**for** $d \leftarrow 1$ **to** $D$ **do**
  Calculate $K(\{d\}, C[i])$ for all non-empty $C[i]$
  **if** $\max(K(\{d\}, C[i]) > t_{BF}$ **then**
    Add $d$ to $C[\arg\max_i K(\{d\}, C[i])$
  **else if** *Not all* $C[1], \ldots, C[N]$ *are empty* **then**
    Add $d$ to first non-empty $C[i]$
  **end**
**end**

**Algorithm 4:** Iterative refinement of clustering assignments

**Result:** $best[1], \ldots, best[D]$ containing the inferred identities
$C[1], \ldots, C[N]$ as exclusive subsets of $\{1, \ldots, D\}$
**for** $iter \leftarrow 1$ **to** $n_{iter}$ **do**
  **for** $d \leftarrow 1$ **to** $D$ **do**
    **for** $i \leftarrow 1$ **to** $N$ **do**
      $L_s[i] \leftarrow \log L_1(d; C[i], \boldsymbol{f})$
      **for** $j \leftarrow 1$ **to** $i - 1$ **do**
        $L_d[i, j] \leftarrow \log L_2(d; C[i], C[j], \boldsymbol{f})$
      **end**
    **end**
    remove $d$ from the current cluster if assigned
    **if** $\max_i L_s[i] > \max_{i,j} L_d[i, j] + t_{singlet}$ **then**
      add $d$ to $C[\arg\max_i L_s[i]]$
      $best[d] \leftarrow singlet(i)$
    **else if** $\max_{i,j} L_d[i, j] > \max_i L_s[i] + t_{doublet}$ **then**
      $best[d] \leftarrow doublet(i, j)$
    **else**
      $best[d] \leftarrow ambiguous$
    **end**
  **end**
  Terminate early if *best* is unchanged.
**end**

### Modeling likelihoods of a single droplet

Let $s \in \{1, 2, \ldots, S\}$ be a sample index, $d \in \{1, 2, \ldots, D\}$ an index of barcoded droplets, and $v \in \{1, 2, \ldots, V\}$ be the index of exonic variants considered (typically MAF $> 1\%$). Let $r_{dv} \in \{0, 1, \ldots\}$, be the read depth overlapping with $v$ from droplet $d$, and let $b_{dvi} \in \{0, 1, 2\}$ be the base call of i-th sequence read ($i \in \{1, 2, \ldots, d_{dv}\}$) consistent to the reference allele (0), the alternate allele (1), or other alleles (2), and $q_{dvi} \in \{0, 1, \ldots\}$ be the phred-scale base quality score[51]. Let $g_v \in \{0, 1, 2\}$ and $e_{dvi} \in \{0, 1\}$ be latent variables representing true underlying genotypes and the event of a sequencing error, respectively. Then $\Pr(b_{dvi} | g_v, e_{dvi})$ is assumed to follow the distribution widely used in other studies[52], where $\Pr(e_{dvi})$ follows Bernoulli $\left(10^{-0.1 q_{dvi}}\right)$.. Then, the probability of allelic read given genotype can be written as $\Pr(b_{dvi} | g_{sv}) = \sum_{e_{dvi}=0}^{1} \Pr(b_{dvi} | g_{sv}, e_{dvi}) \Pr(e_{dvi})$.

Under the assumption that all the reads in barcoded droplet $d$ were originated from one sample (i.e. $d$ is a singlet), the likelihood of singlet can be modeled as follows.

$$L_1 = \prod_{v=1}^{V} \left[ \sum_{g_v=0}^{2} \left( \prod_{i=1}^{r_{dv}} \Pr(b_{dvi}|g_v) \right) \Pr(g_v) \right]$$

where $\Pr(g_v)$ is the probability of unobserved genotype. In demuxlet[11], we assumed that $Pr(g_v)$ is given based on the external genotypes (based on the posterior probability of imputed genotypes or best-guess genotypes with predefined error rates) for each sample. In freemuxlet, we model $\Pr(g_v)$ in multiple ways across different steps. Initially, when the identity of each barcoded droplet is unknown, we assume that $\Pr(g_v)$ only depends on the non-reference allele frequency $f_v \in (0, 1)$, obtained from external resources such as the 1000 Genomes Project. In this case $\Pr(g_v) = \Pr(g_v; f_v) \sim \text{Binomial}(2, f_v)$. Alternatively, $\Pr(g_v)$ can be defined assuming a cluster of droplets $C \subseteq \{1, \ldots, D\}$ contains the sequence reads from the originating sample using genotype likelihoods and Bayes' rule.

$$\Pr(g_v; C, f_v) = \frac{\left(\prod_{d \in C} \prod_{i=1}^{r_{dv}} \Pr(b_{dvi}|g_v)\right) \Pr_{f_v}(g_v; f_v)}{\sum_{g=0}^{2} \left[ \left(\prod_{d \in C} \prod_{i=1}^{r_{dv}} \Pr(b_{dvi}|g)\right) \Pr(g; f_v) \right]}$$

$L_1$ can be defined without cluster (i.e. $L_1(d; \boldsymbol{f})$) and with cluster (i.e. $L_1(d; C, \boldsymbol{f})$), where $\boldsymbol{f}$ is a vector of allele frequencies across all variants. Under the assumption that the reads in $c$ were originated from two samples with mixing proportion $(1 - \alpha) : \alpha$, the likelihood of doublet becomes

$$L_2 = \prod_{v=1}^{V} \left[ \sum_{g_{v1}=0}^{2} \sum_{g_{v2}=0}^{2} \left( \prod_{i=1}^{r_{dv}} \Pr(b_{dvi}|g_v) \right) \Pr(g_{v1}) \Pr(g_{v2}) \right]$$

where $\Pr(b_{dvi}|g_{v1}, g_{v1}; \alpha) = (1 - \alpha) \Pr(b_{dvi}|g_{v1}) + \alpha \Pr(b_{dvi}|g_{v2})$. $\Pr(g_{v1})$ and $\Pr(g_{v2})$ can also be defined with or without cluster, and $L_2$ can be defined without clusters (i.e. $L_2(d; \alpha, \boldsymbol{f})$) and with clusters (i.e. $L_2(d; \alpha, C_1, C_2, f)$) where $C_1$ and $C_2$ are exclusive subsets of $\{1, \ldots, D\}$.

#### Initializing cluster assignments with singlet scores

To initialize our cluster assignment, we define "Singlet Score" of each barcoded droplet ($S(d)$) as a log Bayes Factor between the singlet and doublet likelihood based on allele frequencies only. To compute singlet score, we assume that non-reference allele frequency of each variant is known as $f_v$, and assume that $\Pr(g_v) = \Pr_{f_v}(g_v) \sim \text{Binomial}(2, f_v)$. In this way, we can model likelihoods of singlets and doublets without requiring individual genotypes or clustered droplets, just by modeling unobserved genotypes as latent variables modeled by allele frequencies.

$$S(d) = \log \left[ \frac{L_1(d; \boldsymbol{f})}{L_2(d; \alpha = 0.5, \boldsymbol{f})} \right]$$

The Singlet Score informs whether the scRNA-seq reads from a barcoded droplet is likely singlets or doublets without requiring external genotypes. The likelihood models used to obtain single score is identical to the models used for detecting sample contamination from DNA sequence reads[52], except that $\alpha$ is fixed to 0.5. For droplets with relatively few reads, $S(d)$ may not be much informative and the value will be close to zero. For droplets with larger read counts, $S(d)$ will become more informative and will likely diverge from zero into either direction. When performing clustering, we sort barcoded droplets based on decreasing orders of $S_c$ so that putative singlets are assigned to a cluster first, so that doublets will have less chance to confound the clustering results.

#### Genetic similarity between groups of barcoded droplets

To assign individual droplets to a cluster, freemuxlet utilizes "Genetic Similarity" (K) between two groups of barcoded droplets. Each group may consist of a single droplet or multiple droplets. Let $C_1$ and $C_2$ be exclusive subsets of barcoded droplets $\{1, \ldots, D\}$. Let $C_u \in C_1 \cup C_2$. If the reads from the two groups of droplets $C_1$ and $C_2$ originated from the same individual, its likelihood can be approximated by incorporating $\Pr(g_v; C_u, f_v)$ into the equation.

$$L_3(C_u; \boldsymbol{f}) = \prod_{v=1}^{V} \left[ \sum_{g_v=0}^{2} \left( \prod_{d \in C_u} \prod_{i=1}^{r_{dv}} \Pr(b_{dvi}|g_v) \right) \Pr(g_v; C_u, f_v) \right]$$

On the other hand, if $C_1$ and $C_2$ were originated from two different individuals, the likelihood can be approximated in a similar manner, but allowing two latent genotypes for each cluster.

$$L_4(C_1, C_2; \boldsymbol{f}) = \prod_{v=1}^{V} \left[ \left\{ \sum_{g_1=0}^{2} \left( \prod_{d \in C_1} \prod_{i=1}^{r_{dv}} \Pr(b_{dvi}|g_1) \right) \Pr(g_1; C_1, f_v) \right\} \left\{ \sum_{g_2=0}^{2} \left( \prod_{d \in C_2} \prod_{i=1}^{r_{dv}} \Pr(b_{dvi}|g_2) \right) \Pr(g_2; C_2, f_v) \right\} \right]$$

Note that both $L_3$ and $L_4$ assume that $C_1$ and $C_2$ are singlets, and ignore the possibilities that they are doublets. The genetic similarity between the two the joint likelihood of observed data in two groups of droplets can be defined as the following log Bayes Factor:

$$K(C_1, C_2) = \log \left[ \frac{L_3(C_1 \cup C_2; \boldsymbol{f})}{L_4(C_1, C_2; \boldsymbol{f})} \right]$$

If $C_1$ and $C_2$ contain droplets from the same individual $K(C_1, C_2)$ is expected to have positive values. If they originated from different individuals, $K(C_1, C_2)$ is expected to have negative values. If droplets in $C_1$ and $C_2$ have very few reads, the value will be close to zero. We use this metric as surrogate of similarity/distance when clustering individual droplets.

### PBMC Processing

#### Isolation and Cryopreservation

Informed consent was obtained from all donors sequenced in this study. Frozen PBMCs were obtained from 54 healthy donors and processed using published protocols from the Immune Variation project (ImmVar) [53–55]. These samples were divided into two shipments: "old_immvar" and "new_immvar". PBMCs were also collected from an additional 10 healthy donors by the UCSF AIDS Specimen Bank (ASB) via density gradient centrifugation using Ficoll-hypaque (density 1.077), after which they were washed, counted on an automated cell counter (Beckman Coulter), and aliquoted in cryopreservation media (90% fetal calf serum and 10% DMSO) at 3 million cells/mL for freezing at -80°C. To increase the statistical power of genetic analyses, we restricted the demographics of these cohorts to only female, non-Hispanic white donors, ranging in age from 20 to 56.

#### Processing and Stimulation

For the AMO experiment, cryovials were thawed by swirling in a 37°C water until only small ice crystals remained (approx. 1.5 min). Cells were then gently pipetted and directly deposited into 4.5 mL of warmed media. Cells were counted using a Countess automated cell counter and plated in a 96-well flat-bottom plate at 1 million cells in 200 μL/well as is or with the 3 stimulants (IFN-β, 100 IU/mL; IFN-γ, 100 IU/mL; PMA/I at 1x (81 nM PMA1.33  M Ionomycin)). The plate incubated at 37°C/5%$CO_2$ for 8 hours. Several wells were counted for representative cell recovery values over the incubation, which were then used to withdraw approximately equal cell numbers from each well and mix them into 5 pools according to the design. Cells were washed once with PBS + 0.4% BSA, and counted. Each pool was then split in two aliquots for the separate RNA and ATAC assays. The RNA-seq aliquots were loaded into 5 lanes the 10x Chromium controller using the 3'v2 kit at 50,000 cells per droplet reaction using standard protocols. For the ATAC-seq aliquots, cells were lysed to isolate nuclei according to the demonstrated protocol published by 10x Genomics (document CG000169, Rev D), and then assayed according to the published protocol.

For the production experiment with 64 donors, vials were thawed in 2 batches of 32 vials each in the same manner as the AMO experiment. Each sample was diluted in PBS + 500 nM propidium iodide and CountBright counting beads (Invitrogen, C36950) and flowed on an Attune NXT flow cytometer for counting using the plate reader. Cells from each donor were then plated in four (4) 96-well plates, 6 wells per patient, at $3 \times 10^5$ cells/well. Solutions of each agonists (and PBS control) were added at 4 μL/well to achieve target final concentrations (same as AMO, and with TNF-α, 50 ng/mL; R-848, 1 μg/mL). Cells incubated for 9 hours at 37°C and 5% CO2. After incubation, plates were placed on ice and samples were taken for counting using the cytometer as described above. To avoid counting all 384 wells, samples from only 99 wells were taken: the control well from each of 64 individuals + 35 agonist wells, 7 each of 5 conditions, from random individuals. We confirmed there was no prominent count differences due to condition, and extrapolated the control count of each individual to all of that individual's samples to determine volumes for pooling. Cells from each individual were pooled into 12 tubes according to the experimental design with the plates on ice. Cells pre-incubated with Fc block (Biolegend) for 10 minutes on ice and then stained with a universal cocktail of 99 antibody-oligonucleotide conjugates (BD Genomics), washed 2x with of PBS + 0.4% BSA, strained with a 40 um strainer. All 12 pools were then loaded onto the 10x Chromium instrument, one per lane (droplet reaction).

### FASTQ Processing and Demultiplexing

#### Alignment and Counting

Bulk RNA-seq data was aligned and counted according to the QuantSeq 3' mRNA-Seq Integrated Data Analysis Pipeline (Lexogen), using STAR alignment to the GRCh38-3.0.0 transcriptome reference (10x Genomics) and HTSeq v0.6.0. Single-cell transcriptome data were aligned and counted using cellranger (10x Genomics), v3.0.2 for the AMO experiment and v3.1.0 for the production run. with the GRCh38-3.0.0 transcriptome reference. Single-cell antibody-derived tags (ADT) data were aligned and counted using cellranger v3.1.0 with a custom reference. Single-cell ATAC-seq data were aligned and counted using cellranger-atac v1.2.0 with genomic reference GRCh38-1.1.0.

#### Freemuxlet and Decoding

The freemuxlet algorithm, as implemented as a part of the popscle suite of population-scale analysis tools for single-cell genomics, involves pre-processing the binary sequence alignment files (BAMs) output by cellranger using dsc-pileup, a sub-command which annotates the base calls at specified sites throughout the reference genome. To generate the list of sites, we used the variant calling format (VCF) file provided by the 1000 Genomes Project (1KG) ALL.wgs.shapeit2_integrated_snvindels_v2a.GRCh38.27022019.sites.vcf.gz, using only autosomal sites classified as "common", called with high variant confidence (QD > 10.0), with a minor allele frequency (MAF) of 0.01 and minor allele count (MAC) of 1. For RNA-seq data, we additionally filtered the 1KG VCF for exonic sites using UCSC annotations. For ATAC-seq data in the AMO experiment, we additionally filtered by intersecting the peaks.bed files output by each of the 5 cellranger-atac outputs with the pre-processed 1KG VCF. The VCF, BAM, and filtered whitelist of cell barcodes were fed into dsc-pileup, to generate one set of pileup files per droplet reaction (pool).

In order to allow the clustering algorithm to leverage the genetic droplet data across multiple reactions, we merged the sets of pileup files (5 for AMO, 12 for production run) using a custom script, changing the GEM group to avoid cell barcode collisions. We then fed the single merged set of pileup files into the freemuxlet sub-command of popscle, with –nsample 5 for AMO and –nsample 64 for production run, and with the –aux-files parameter set to have the pairwise Bayes factor for each possible pair of droplets reported in the final output ("ldist" file). Freemuxlet was run on a cloud compute instance (AWS, m5a.24xlarge) running Ubuntu 18.04 with 96 vCPUs and 384 GiB of memory, and completed in approximately 128 hours. The freemuxlet implementation of popscle has since been improved for efficiency and memory allocation.

#### Validation of Freemuxlet Genotypes

ImmVar samples were genotyped in two batches, both using the OmniExpress Exome SNP array (Illumina), while ASB samples were genotyped using the World LAT Array (Affymetrix). Raw files from each of the genotyping arrays were converted into 3 separate VCFs using PLINK v1.90. Using a sample subset of the ImmVar VCF containing only the 5 donors in the AMO experiment, demuxlet was run separately on each of the 5 output BAM with a whitelist of cell barcodes and default parameters. Genotype similarity between pairwise VCF sample IDs was computed as described above, implemented for VCFs using a custom script (https://github.com/hyunminkang/apigenome/blob/master/scripts/vcf-match-sample-ids). To generate the visualizations of genetic distance, the "ldist" file (one pair of droplets per line) was reshaped into a pairwise distance matrix, corrected by adding a constant to make distances non-negative, and input into Python UMAP v0.4.6 with parameter metric="precomputed".

### Single-cell Data Processing and Analysis

#### Pre-processing and Low Quality Cell Removal

For each experiment, individual transcriptome cellranger outputs from each of the 12 droplet reactions were aggregated using cellranger aggr to create a single counts matrix. Decoding of pooling matrix was done as described above, and delineated in Figure S5A, to map the anonymous freemuxlet clusters to the known donors and attach demographic information and experimental covariates (i.e. viability, stimulation condition). All analyses that follow were done with Python v3.6.10 and Scanpy v1.5.1. To create an initial UMAP visualization, the genes were filtered to a minimum count of 100. Counts were then normalized per cell (to a sum of 1e6 per cell), log1p transformed, and scaled to mean 0 and variance 1. The ComBat algorithm was then run to correct for batch across the 12 reactions, with included covariates for stimulation condition and donor ID. The corrected data was scaled again and 200 principal components were computed. A neighborhood graph was computed using the top 175 principal components with a local neighborhood size of 15. UMAP coordinates (default parameters) and clusters (Leiden algorithm, with resolution=1) were then computed and visualized.

Upon initial visualization, we noted a prominent mitochondrial effect present across all batches, where a bimodal distribution across cells was observed when plotting the percentage of transcripts aligning to mitochondrial genes ("percent mito"). High percent mito in single-cell PBMC data indicates a stress and/or apoptosis response, a decrease in viability we attributed to the extended time required to manually pool and stain cells given the size of the experiment. Phenotyping high-mito clusters suggested a 1-to-1 correspondence between low and high mito clusters. Rather than attempting to correct for this, we decided to exclude the high percent mito cells from further analysis. We also observed some effects due to low counts of either genes or proteins, as well as some clusters marked by platelet and red blood cell genes, and also excluded these cells from further analysis.

Using these re-filtered cell barcodes from the transcriptome data, we re-ran the processing as described from the raw values, now carrying forward 150 PCs, which resulted in the final UMAP visualization shown in the main figures. Surface protein data was then concatenated across wells and joined to the transcriptome data using the re-filtered barcodes. Summing counts per antibody identified 3 antibodies with nearly two orders of magnitude fewer counts than the rest of the 99-antibody panel: CD40, CD44, and CD19. These proteins were excluded from further analysis. The counts data for the remaining 96 antibodies were normalized per cell (to a 1e6 sum per cell) and centered log-ratio transformed.

#### Differential Expression Analysis and Identifying Cell-type-specific transcriptional Responses to Perturbations

Differential expression analysis was performed using R v3.6.1 and DESeq2 v1.26.0. Raw gene and protein expression counts were pseudobulked (summed) per donor-cell type-condition, at different resolution levels of cell type annotation. For PMA/I-stimulated myeloid cells that did not cluster out into individual myeloid subtypes, corresponding pseudobulks were generated for the other stimulation conditions for comparison purposes. Stimulation conditions were then compared to control for each cell type, with formula $\sim condition$. For the interferon analysis, the interferon conditions were directly compared to each other using the same formula.

To generate the heatmap in Figure 3F, we extracted out differentially expressed genes using cutoffs of $abs(log_2(FC)) > 1.5$ and $p_{adj} < 0.05$. From that list, in an effort to balance the number of DE genes per sample, we used only the top 200 significant genes from each comparison, since some stimulation conditions had 1000s, while others has only 10s or low 100s. Taking the union of these filtered gene sets resulted in 1853 genes across 62 cell-type-condition groups, which were then $k$-means clustered into gene modules with $k = 40$. Mean $log_2(FC)$ was computed and plotted in the heatmap for each cell-type-condition-gene module, and rows and columns were ordered using hierarchical clustering with Scipy v1.5.3 scipy.cluster.hierarchy, with parameters "method=average" and "optimal_ordering=True". Gene modules were then identified using ToppGene functional enrichment (https://toppgene.cchmc.org/enrichment.jsp) and manually annotated using the significant Gene Ontology terms.

To identify cell-type-specific transcriptional responses to perturbation as plotted in Figure S8A, we extracted out differentially expressed genes using cutoffs $log_2(FC) > 0.5$ and $p_{adj} < 0.1$, the more liberal cutoffs being used in an effort to identify genes that may be only modestly upregulated but highly specific. For a given gene $g$, stimulation condition $c$, and cell type $t$, we calculate the mean expression across all other cell types, $\mu_{g,c,t_o}$. We then calculate a ratio, $r_{g,c,t}$:

$$r_{g,c,t} = \frac{log_2(FC_{control})_{g,c,t}}{\mu_{g,c,t_o} + 1}$$

We use these ratios to rank the genes and identify those whose upregulation is highly-specific to cell type and perturbation.

#### BiNGO and Gene Ontology Analysis using Cytoscape

To generate Gene Ontology network graphs in Figure 5B, we used BiNGO v3.0.5 and Cytoscape v3.9.1 running Java v11.0.6. Using the DESeq2 genes identified with $log_2(FC) > 0$ and $p_{adj} < 0.05$, we generated an upregulated list of genes for each of the interferons IFN-$\gamma$ and IFN-$\beta$. We used this list to perform a hypergeometric test to assess over-representation, with a Benjamani-Hochberg false discovery rate (FDR) of 0.05, using the release of the go.obo file accessed from geneontology.org on June 8, 2022, with the whole annotation as a reference set and using only the "biological process" namespace for Homo Sapiens. After computing the network graph, the coordinates were exported from Cytoscape and loaded into the Python environment. The adjacency matrix from the network graph was then fed as the connectivity neighbor graph into Leiden clustering (resolution: cM, 1.2; ncM, 1.0) to identify "pathway clusters" whose ontologies were related based on shared genes.

### Genetics and eQTL Analysis

#### Imputation and eQTL Analysis using MatrixeQTL

Imputation was performed separately for the 3 VCFs (ImmVar-1, 55 individuals; ImmVar-2, 27 individuals; and ASB, 10 individuals). Samples were first evaluated for missingness, Hardy-Weinberg equilibrium, and heterozygosity with Plink, and 558,534 (ImmVar-1), 564,069 (Immvar-2), and 501,587 (ASB) SNPs were retained for further analysis. We then performed imputation using the Michigan Imputation Server's web portal with the Haplotype Reference Consortium version 1.1 reference set. The datasets were further filtered to 557,262 (ImmVar-1), 562,460 (ImmVar-2), and 494,667 (ASB), SNPs when accounting for invalid alleles, multi allelic sites, monomorphic sites, allele mismatches, and SNP call rates less than 90%. A total of 9,652,826 (ImmVar-1), 8,737,184 (ImmVar-2), and 6,434,802 (ASB) SNPs were imputed with an $R^2 > 0.3$. Imputed SNPs from the three cohorts were merged and further filtered to only those with a minor allele frequency >10% that were imputed in all individuals, resulting in 4,208,073 SNPs.

Approximately 4.2 million SNPs with a MAF > 10% were used to map *cis*-eQTLs within a window +/- 100kb of each gene. All genes with at least 50 UMI counts in 20% of samples were retained, resulting in a total of 6,522 genes tested. Pseudobulk gene expression profiles were created for each cell type by averaging the single-cell read counts and applying log-normalization. *Cis*-eQTLs were mapped separately for each cell type using the MatrixEQTL R package. The first 5 principal components of gene expression, first 3 genotype principal components, age, and SNP array were included as covariates in all the eQTL linear models. Multiple testing correction was performed using the Benjamini-Hochberg (BH) procedure.

To generate the cell type Manhattan plots in Figure 6A, significant (BH < 0.05) eQTLs were color coded by condition and plotted over downsampled (5%) insignificant eQTLs, colored grey. To make the condition-colored lineplots along the x axis, median $-log_{10}(p)$ was calculated per 25 million base pair bin tiled along the genome, followed by a cubic 1D interpolation for smoothing. To generate the effect size scatter plots in Figure 6D, eQTLs were filtered for the most significant eQTL per gene-cell type-condition to ensure each gene only appears once in the plot. Effect size ($\beta$) was transformed to $\beta_t$ using the following:

$$p = 3$$

$$\beta_t = log_{10}(\beta + 10^{-p}) + p$$

This monotonic transformation yielded a roughly symmetric plot along the diagonal with minimal skewing which made the effect sizes easier to visualize and compare across conditions.

In Figure 6E and 6F, a zscore threshold of ±2.5 (6E) and ±3.0 (6F) was used to filter data points to better visualize the box and whisker plots.

#### eQTL-ATAC Enrichment Analysis

To generate the enrichment box plot in Figure 6B, ATAC peaks were called for all control cells from the AMO experiment using Macs2 v2.2.7.1, with parameters shift=75, extsize=150, nomdel, call-summits, nolambda, keep-dup=all, and q=0.05, then intersected with significant eQTLs (per cell type) using pybedtools v0.8.1 (backend bedtools v2.28.0). The percentage of insignificant eQTLs was calculated from an average of 10 trials per cell type, randomly sampling a matched number of eQTLs from the matrixeQTL output.

To generate the enrichment box plot in Figure 6C, differentially-accessible peak sets between the 4 listed cell types were identified using ArchR v0.9.5, function getMarkerFeatures, where a $log_2(FC)$ and $FDR$ is reported per cell type. Form those outputs, we filtered to only those significant peaks ($FDR < 0.05$) in the top 50% of $log_2(FC)$ per cell type. While ArchR's default parameters report a 500 bp peak, which works well for LSI and dimensionality reduction, for enrichment analysis we widened each peak by 2 KB on either side (4.5 KB peak), and used those to intersect with the reported eQTLs and perform a MannWhitney U test to determine whether reported eQTL p values were higher within peaks than without. The test was run pairwise between eQTL cell types and ATAC cell types, and the $-log_{10}p$ of each test was plotted.
